# Repeated trends in altitudinal gradients of diversity: how habitat filtering and biotic interactions structure ecological communities

**DOI:** 10.64898/2026.03.18.712335

**Authors:** Raphaël Fougeray, Arlety Roy, Christelle Penager, Guillaume Correa Pimpao, Ronald Mori Pezo, Léo-Paul Charlet, Nino Page, Ombeline Sculfort, Stéphanie Gallusser, Marianne Elias, Melanie McClure

**Affiliations:** Laboratoire Écologie, Évolution, Interactions des Systèmes Amazoniens (LEEISA), Université de Guyane, CNRS, IFREMER, 97300 Cayenne, France; Universidad Nacional Autónoma de Alto Amazonas, Yurimaguas, Loreto, Perú; Independent entomologist, French Guiana, France; Instituto de Investigación Biológica de las Cordilleras Orientales, Tarapoto, Perú; Institut de Systématique, Evolution, Biodiversité, Muséum National d’Histoire Naturelle, CNRS, Sorbonne-Université, EPHE, Université des Antilles, Paris, France; Centre Interdisciplinaire de Recherche en Biologie, Collège de France-CNRS-INSERM, Paris, France; Smithsonian Tropical Research Institute, Gamboa, Panama

**Keywords:** Community assembly, Ithomiini, Müllerian mimicry, mutualism, Neotropics, phylogenetic structure, predation, regional species pool

## Abstract

Understanding how biodiversity is structured along tropical elevational gradients requires disentangling the relative roles of regional evolutionary history and local processes shaping ecological assemblies. Here, Ithomiini butterfly communities were studied along repeated elevational gradients in two Neotropical regions with contrasting evolutionary histories: the Amazonian Andes and the Guiana Shield. The study tested whether similar elevational patterns of taxonomic, mimetic, and phylogenetic structure emerge despite distinct regional species pools, and whether abiotic and biotic factors contribute to shaping these patterns.

Despite marked regional differences in overall richness, consistent elevational patterns emerged across both regions. Taxonomic and mimetic richness increased with elevation and were accompanied by stronger phylogenetic clustering, indicating that similar habitat filtering processes operate along altitudinal gradients irrespective of regional context. Phylogenetic β-diversity was predominantly driven by lineage turnover, particularly in the Andes, highlighting the role of elevational gradients in promoting replacement of phylogenetically distinct lineages rather than simple species loss. These shared patterns suggest that altitude has a strong and repeatable effect on community structure, with habitat filtering acting locally on regionally distinct species pool.

Abiotic factors such as temperature appeared to constrain species distributions at broad spatial scales, whereas biotic interactions acted more locally. In particular, butterfly diversity was positively associated with potential host plant richness and predation pressure, indicating that ecological interactions can further shape local community composition once broad-scale environmental constraints are accounted for. By integrating phylogenetic structure, biotic interactions, and environmental gradients across regions with contrasting evolutionary histories, this study shows how regional species pools and local ecological filtering jointly shape tropical biodiversity and highlights that similar elevational assembly processes could arise independently across the Neotropics.

## Introduction

A central goal of ecological research is to understand geographic patterns of species diversity and coexistence, both for its theoretical importance and for setting conservation priorities. One of the best-known patterns of species diversity is the latitudinal gradient of increasing species richness from the pole towards the equator (Hillebrand, 2004). A parallel trend has been observed along altitudinal gradients, where species richness generally decreases with increasing elevation, often exhibiting a hump-shaped pattern with maximum richness at intermediate elevations. Although elevational gradients have been increasingly used to investigate biodiversity patterns and underlying mechanisms, they remain less comprehensively documented and generalized across taxa and regions than latitudinal gradients, with considerable variability among systems (Rahbek, 1995; McCain & Grytnes, 2010; Sundqvist et al., 2013; Quintero & Jetz, 2018). Many of the hypotheses exploring these diversity gradients are interdependent, and current research suggests that in many cases multiple aspects must be considered to fully understand the observed patterns. Such hypotheses can be split into three general groups: 1) historical and evolutionary hypotheses that focus on the duration of an environment and rates of diversification (Fischer, 1960; Pyron & Wiens, 2013), 2) spatial and climatic hypotheses that focus on the extent of the environments and habitat filtering (Rosenzweig, 1995; Jansson & Dynesius, 2002), and 3) ecological hypotheses that focus on mechanisms of species specialization and coexistence (Janz et al., 2006; Forister et al., 2015).

Changes in abiotic and biotic conditions take place over short spatial distances along altitudinal gradients. Repeated studies along these gradients can be useful to disentangle the effects of individual factors on biodiversity, which are often confounded by their natural covariance. However, patterns observed locally emerge from the interplay between processes operating at different spatial and temporal scales, notably the composition of the regional species pools and the factors responsible for local community assemblages (Zobel, 2016; Chase & Myers, 2011). Regional species pools, shaped by historical and evolutionary processes such as diversification, dispersal, and extinction, define the set of species potentially available for local assemblages, whereas local abiotic filtering and biotic interactions determine which species persist under given environmental conditions (Cornell & Harrison, 2014; Mittelbach & Schemske, 2015).

This multi-scale perspective is particularly relevant in tropical montane systems, where steep environmental gradients coincide with strong historical and biogeographic contrasts. A comparative approach of a given biological system across regions enables us to assess whether similar local assembly processes generate convergent biodiversity patterns despite differences in regional species pools. In this context, Ithomiini butterflies are an especially suitable model system. This highly diverse Neotropical tribe (Beccaloni, 1997; Chazot et al., 2019; Doré et al., 2023), originated in the Andes, where extensive diversification occurred, before colonizing adjacent and then more distant lowland regions, with subsequent upslope colonization occurring from lowland source populations (Chazot et al., 2019). Such a biogeographic history implies marked differences in regional species pools beyond differences in species richness, with Andean assemblages reflecting longer diversification times and potentially higher phylogenetic turnover than assemblages in younger regions such as the Guiana Shield. In the Andes, Ithomiini communities exhibit clear altitudinal patterns, with increasing phylogenetic clustering at higher elevations (Chazot et al., 2014; De-Silva et al., 2016). These patterns likely reflect habitat filtering driven by both abiotic and biotic factors that vary predictably with elevation, and may be observed in other tropical montane regions. However, the relative contribution of these ecological factors in shaping phylogenetic structure remains largely unresolved.

In particular, temperature and precipitation may directly shape species distributions through physiological constraints (Montejo-Kovacevich et al., 2020), and indirectly through variation in host plant availability along altitudinal gradients (Gentry, 1988). Because Ithomiini butterflies exhibit strong host plant specialization (primarily on Solanaceae: Willmott & Freitas, 2006), they likely experience strong inter- and intra-specific competition when resources are limited (Kaplan & Denno, 2007). Consequently, regions of high host plant diversity may offer increased ecological opportunities, facilitating host shifts and thus promoting diversification—a pattern consistent with the hypothesis that diversity begets diversity (Janz et al., 2006; Stevens & Tello, 2011). Furthermore, aposematism in Ithomiini (warning coloration signalling unpalatability: Poulton, 1890) and their numerical dominance within mimicry rings (co-occurring species sharing similar warning signals: Beccaloni 1997) make them especially sensitive to predation-driven selective pressures. Predator-mediated selection promotes mutualistic convergence in warning signals (Müller, 1879), reinforcing mimicry systems and influencing community assembly by shaping which phenotypes and lineages can coexist locally (Elias et al., 2008; Chazot et al., 2014). Through this process, predation can facilitate species coexistence via microhabitat partitioning and ecological specialization, while simultaneously constraining phenotypic divergence through selection toward shared warning signals (Willmott et al., 2017; Aubier & Elias, 2020). As a result, spatial variation in predation pressure across multiple spatial scales may generate differences in the selective environments experienced by visually conspicuous butterflies, with potential consequences for trait evolution as well as the taxonomic, mimetic, and phylogenetic structure of communities along environmental gradients. Collectively, these ecological characteristics position Ithomiini butterflies as an ideal model for examining the interplay between biotic interactions and habitat filtering in shaping biodiversity patterns across distinct altitudinal and geographical contexts.

Despite substantial research on altitudinal biodiversity patterns, efforts to reconcile the relative importance of abiotic constraints and biotic interactions have yielded results that remain highly context- and system-dependent (Sundqvist et al., 2013). This variability likely reflects differences in regional histories (Kraft et al., 2011), environmental heterogeneity (Stein et al., 2014), and the spatial scales at which community assembly processes operate (Cornell & Harrison, 2014). In particular, comparative studies across biogeographic regions remain relatively scarce (but see Kessler et al., 2011 and Myers et al., 2013) and rarely integrate effect on phylogenetic structure. This study addresses this gap by examining the relative and interactive roles of abiotic factors (temperature, precipitation) and biotic interactions (host plant availability, predation pressure, mimetic interactions) in shaping the structure, diversity and phylogenetic composition of Ithomiini butterfly communities along altitudinal gradients in two distinct regions characterized by contrasting regional species pools—the Amazonian Andes, where Ithomiini originated and diversified over long evolutionary timescales, and the Guiana Shield, which harbours assemblages derived from more recent colonization events (Chazot et al., 2019).

Specifically, we hypothesize that (i) the taxonomic, phylogenetic, and mimetic structure of Ithomiini communities varies with elevation in both regions following similar patterns. Despite the contrasting geological and biogeographic histories of the Andes and the Guiana Shield, such similarities in altitudinal patterns would suggest that comparable ecological and evolutionary processes drive local community structuring across Neotropical montane environments; (ii) community composition strongly correlates with abiotic factors, particularly temperature and precipitation, along elevation gradients, as a result of habitat filtering acting primarily at broader spatial scales through physiological constraints on species distributions (Dewan et al., 2022) and indirectly through effects on host plant availability (Park et al., 2020); (iii) taxonomic, mimetic, and phylogenetic diversity are influenced locally by biotic interactions, notably potential host plant diversity and predation pressure. Greater potential host plant diversity may increase ecological opportunities, promoting butterfly diversification (“diversity begets diversity”), whereas stronger predation pressure, tested here using field-based predation experiments, is expected to constrain variation in aposematic signals, reducing mimetic diversity within communities while simultaneously strengthening mimicry-driven mutualism. By explicitly testing these hypotheses through a comparative regional approach, this study provides novel insights into the generality of ecological and evolutionary processes structuring tropical biodiversity, contributing to our broader understanding of community assembly in complex ecosystems.

## Methods

### Study sites

Sampling was undertaken across 12 sites in two regions: Peru (departments of Loreto and San Martín) and French Guiana, located on the northern coast of South America. Six sites were surveyed in each region during the dry seasons of 2022 and 2023. Sites were distributed across three different altitudinal ranges (2 sites per altitude per region: low (<200m), mid (250-500 m), and high (>700 m); see Supporting Information and map in Fig. 1a for coordinates). Three sites in French Guiana (one from each altitude) were re-sampled during the rain season of 2024 to assess the potential influence of seasonality on community composition and were not included in any other analyses.

**Figure 1.**
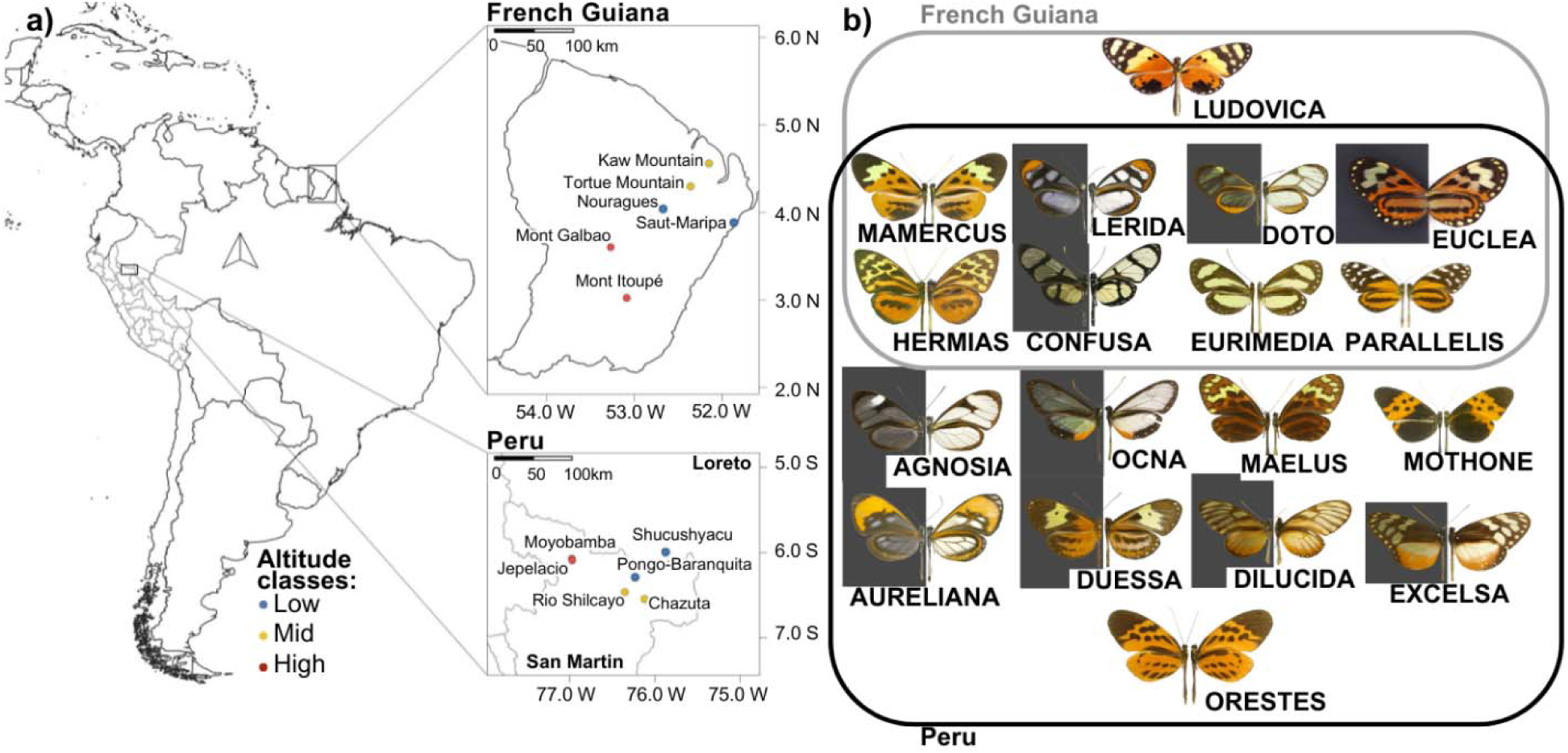
Ithomiini community composition and diversity (taxa and mimicry) in Peru and French Guiana were assessed along an altitudinal gradient. Shown here are a) the map of the study sites, with colours indicating altitude classes, and b) photographs of species (not at scale) representative of the mimicry rings of Ithomiini butterflies observed in this study, shown by region. Dorsal views are on the left (against a dark background for transparent-winged species), and ventral views on the right. All photographs were taken by Keith Willmott, except for EUCLEA and LUDOVICA, which were taken by Raphaël Fougeray.

Sampling was standardized across sites, with three collectors operating simultaneously and continuously from 07:00 to 16:00 for four consecutive days at each site. Sampling consisted of active searches along forest trails and surrounding forest (corresponding to an area of approximately ∼0.5–2 km²) covering a broad range of microhabitats and flight heights typical of Ithomiini butterflies. Collected specimens were sexed, identified (primarily at the subspecies level, referred to as ‘taxa’), and classified into one of 46 mimicry rings (see Fig. 1b); 44 commonly referenced as per Doré et al. (2022), and 2 additional ones (EUCLEA and LUDOVICA) due to their sufficiently distinctive wing pattern and associated mimetic community. Rarefaction curves were used to assess whether sampling effort was sufficient (iNEXT.3D R package; Chao et al., 2021), and as sample coverage reached an asymptote (Fig. S1), was assumed to be adequate for further analyses. No ethical approval was required for this study.

### Spatial patterns of diversity and phylogenetic community structure in Ithomiini

To quantify diversity, richness (*S*), Shannon’s diversity index (*H*), and Pielou’s evenness (*J*), were calculated separately for both taxa (*T*) and mimicry-rings (*MR*) using the R package vegan (Oksanen et al., 2024). To test whether species and mimicry-ring richness differed between French Guiana and Peru, and whether altitudinal patterns in diversity were consistent across regions, generalized linear mixed models (GLMMs) with Poisson distributions were applied for each response variable and separately for taxa richness and mimicry-ring richness. Model assumptions for Poisson GLMMs were validated through inspection of residuals to check for overdispersion. Post-hoc Tukey tests were further used to assess differences between altitudes using the multcomp R package (Bretz et al., 2016).

To test whether Ithomiini communities exhibited similar altitudinal patterns of species and mimicry-ring composition in both regions, we applied permutational multivariate analysis of variance (PERMANOVA, R package vegan) to Bray–Curtis dissimilarity matrices of taxa and mimicry rings between sites, which are appropriate for abundance data and reduce sensitivity to differences in total counts among sites. Post hoc pairwise comparisons were performed using pairwise Adonis R package (Martinez Arbizu, 2020). Tests of multivariate dispersion indicated homogeneous dispersion among altitude classes and regions for both taxa and mimicry-ring community matrices (see Supplementary Information for details). As replication was limited, the significant interaction between altitude and region associated with near-zero residual variance was not considered reliable. Finally, non-metric multidimensional scaling (NMDS) ordinations were used to provide a visualization of multivariate community dissimilarities.

To compare phylogenetic diversity, α- and β-diversity indices were calculated using the most recent phylogeny of the Ithomiini tribe (Chazot et al., 2019). To evaluate whether patterns of phylogenetic clustering with elevation were consistent between both regions, linear mixed models (LMMs) were fitted separately for the net relatedness index (NRI) and the nearest taxon index (NTI) (Webb et al., 2002; see Supplementary Information). Assumptions of normality and homoscedasticity of residuals were validated through diagnostic plots. Post-hoc pairwise comparisons were done using estimated marginal means (R package emmeans: Lenth, 2025), applying Tukey’s correction for multiple testing.

To assess whether lineage composition varied across altitude classes in both regions, phylogenetic β-diversity between sites was quantified using UniFrac distance (Lozupone & Knight, 2005). The UniFrac distance was analyzed using PERMANOVA to assess the effect of altitude and region on phylogenetic β-diversity followed by pairwise comparisons. Homogeneity of multivariate dispersion was confirmed for the main effects of altitude and region, whereas the interaction between altitude and region was not retained due to heterogeneous dispersion (see Supplementary Information). Phylogenetic β-diversity was then partitioned into turnover and nestedness using the R package betapart (Baselga & Orme, 2012; Leprieur et al., 2012). This partitioning allows discrimination between community dissimilarity driven by the replacement of lineages between sites (phylogenetic turnover) and dissimilarity resulting from differences in phylogenetic richness, where one community represents a subset of another (phylogenetic nestedness). This distinction is particularly relevant along altitudinal gradients, as it helps identify whether changes in community composition primarily reflect lineage replacement across environmental conditions or the progressive loss or gain of lineages. To assess whether the relative contribution of phylogenetic turnover and nestedness to community dissimilarity were consistent across regions, a generalized linear model (GLM) with a Tweedie distribution was fitted using the R package cplm (Zhang, 2013; see Supplementary Information). Model assumptions were validated using diagnostic plots and by testing for heteroscedasticity via a Spearman correlation between absolute residuals and fitted values (ρ = −0.16, p = 0.21).

### The role of altitude in habitat filtering

#### Climatic variation and elevation: temperature and precipitation gradients

To record temperature data during the sampling period, three HOBO sensors (HOBO Pendant MX Temperature/Light Data Logger MX2202) were placed at each site for 4 days and were used to obtain the average daily temperature between 06:00 - 19:00. In addition to these field measurements, three independent climatic predictors were also used for each site: average annual temperature and precipitation data from 1970 to 2000 were obtained (with 30” resolution, ≈1km²) from WorldClim (Fick & Hijmans, 2017), and rainfall data for the sampling days were obtained from AgERA 5 (Boogaard et al., 2020). To test whether temperature and precipitation were significantly correlated with elevation, generalized linear models (GLMs) were performed separately for both Peru and French Guiana. For all variables, Gamma GLMs were used after validation of overdispersion through residual diagnostics. Rainfall in Mont Galbao, French Guiana, was an outlier and therefore excluded (see Supplementary Information).

#### Effect of seasonality and short-term climatic conditions on Ithomiini assemblages

A PERMANOVA was used on Bray–Curtis dissimilarity matrices of taxa abundances for the three sites in French Guiana re-sampled during the rain season. Multivariate tests confirmed homogeneous dispersion for the main effects of altitude and season based on Bray–Curtis dissimilarities (see Supplementary Information).

#### Predation pressure across elevation and its influence on diversity and mutualism

To assess predation pressure, 200 artificial butterfly models resembling local, cryptic and palatable species that are locally abundant were used at each site (*Pierella hyceta* in Peru and *Pierella astyoche* in French Guiana - see Fig. S2 and Supplementary Information). Models were placed every 10 meters along transects at varying heights (at approximately 50 cm and 2 m) and left in situ for 3 days in the same sites where butterflies were collected. Avian predation rate was calculated based on the number of models attacked by birds divided by the total number of models retrieved at the end of the experiment (see Fig. S3 for examples).

To test whether predation pressure varied along altitude classes, a binomial generalized linear mixed model (GLMM) was fitted with region as a random effect. Additional GLMMs (Poisson or quasi-Poisson distribution) were performed to test whether predation pressure influenced species richness, mimetic richness, or the median number of species per mimicry ring, as expected per hypotheses of diversity-dependent selection or mutualistic convergence. Assumptions of the binomial, Poisson, and quasi-Poisson GLMMs were verified through residual diagnostics, indicating no evidence of overdispersion.

#### Potential host plant diversity across elevation: the “diversity begets diversity” hypothesis

The relationship between Ithomiini butterflies and potential host plant species richness was tested in French Guiana, where more abundant and reliable plant occurrence data were available. Potential hosts were defined based on well-established larval host associations documented for Ithomiini butterflies, which predominantly feed on Solanaceae and Gesneriaceae (Willmott & Freitas, 2006). Occurrence data for potential host plants (3,055 occurrences, 88 species) were sourced from the global biodiversity information facility (GBIF) and records from the French Guiana IRD (French institute for development) herbarium. Ithomiini occurrence data (1684 occurrences, 38 species) were compiled from the wildlife observation database of French Guiana Faune-Guyane (GEPOG, 2024), our own field sampling, and grid-cell distribution records from Doré et al. (2022) and Willmott et al. (2021). Species richness was modelled using ensemble species distribution models (SSDMs) with target-group backgrounds to correct for sampling biases (Elith et al., 2010; Barber et al., 2022; Fig. S4) and climatic predictors from WorldClim (Fick & Hijmans, 2017; see Supplementary Information).

To test whether potential host plant richness was positively correlated with Ithomiini species richness, as predicted under the “diversity begets diversity” hypothesis, a generalized additive model (GAM) with a quasi-Poisson distribution was fitted. This allowed for the possibility of non-linear relationships between butterfly and potential host plant richness along the elevational gradient. The analysis was based on data extracted from SSDMs, with each data point representing a pixel from which predicted values of potential host plant richness and Ithomiini richness were retrieved and used as input variables in the model. The model showed no issues with dispersion, residuals, or smoothing, and performance was evaluated using the proportion of deviance explained and F-statistic. To further illustrate how Ithomiini and potential host plant richness were structured along the elevational gradient, ordination-based analyses were used for visualization (see Supplementary Information).

#### Ecological predictors of Ithomiini assemblages

To assess whether Ithomiini communities were influenced by ecological conditions, mixed multivariate distance matrix regression (mixed MDMR package in R) were used (McArtor et al., 2016). All ecological variables were standardized to ensure comparability and tested for multicollinearity. The retained environmental variables included annual mean temperature (WorldClim), annual precipitation (WorldClim), predation rate and mean forest cover (based on five photographs per site; see Supplementary Information). This multivariate approach models variation in community dissimilarities (Bray–Curtis) as a function of multiple predictors simultaneously, while allowing the incorporation of random effects. In contrast to distance-based redundancy analysis (db-RDA), mixed MDMR can account for region as a random factor, making it better suited for comparative analyses across biogeographic regions (Zapala & Schork, 2012).

## Results

### Spatial patterns of diversity and phylogenetic community structure in Ithomiini: the effect of evolutionary history does not override the influence of altitude

A total of 1967 Ithomiini butterflies (96 taxa, 29 genera, and 18 of the 44 potential mimicry rings) were collected, of which 1474 individuals (76 taxa and 27 genera, 16 mimicry rings) were sampled in Peru and 493 (29 taxa and 11 genera, 9 mimicry rings) in French Guiana, plus an additional 225 individuals (30 taxa and 11 genera, 9 mimicry rings) collected in French Guiana during the rain season. Inter-regional comparisons between Peru and French Guiana revealed significant differences in richness (Table 1). Taxa richness (*S_T_*) and mimicry richness (*S_MR_*) were significantly higher in Peru than in French Guiana (*S_T_*: β = 15.8, p < 0.001; *S_MR_*: β = 4.5, p = 0.006) with an increase in taxa richness at higher altitudes (*S_T_*: low-mid: β = 2.42, p = 0.655; low-high: β = 9.64, p = 0.005; mid-high: β = 7.22, p = 0.054). A similar, though non-significant, trend was observed for mimicry richness between low and high altitude (*S_MR_*: low-high: β = 1.81, p = 0.63). Species diversity, as measured by the Shannon index (*H*), and Pielou’s evenness index (*J*) showed similar patterns between Peru and French Guiana, with a decrease in low-altitude sites. PERMANOVA showed that altitude was a key driver of taxa (*T*) and mimicry-rings (*MR*) communities, explaining, for both, 19% of the total variance (*T*: p = 0.016, MR: p = 0.039), with significant differences between high and low altitudes for both taxa and mimicry ring communities (*T*: R² = 0.10, p = 0.037; *MR*: R² =0.20, p = 0.02), whereas differences with mid-altitudes were not significant (p > 0.05), except for differences in mimetic communities between low and mid-altitudes (R² = 0.19, p = 0.04). This can be seen in the NMDS plots (Fig. S5), which shows clear clustering with altitude, although clustering is less apparent at mid-altitudes.

**Table 1.** Diversity indices by study site. Including species richness *S*, the Shannon index *H*, Pielou’s evenness index *J*, the average number of taxa per mimicry-rings μ*TMR* and the median number of taxa per mimicry-rings *MeTMR*. Subscript *T* refer to taxa, while *MR* refers to mimicry-rings. The sites are ordered by decreasing altitude, and the altitude class and altitude for each site is indicated in parentheses.

| Region | Sites (altitude class - altitude) | $S_T$ | $H_T$ | $J_T$ | $S_{MR}$ | $H_{MR}$ | $J_{MR}$ | $\mu TMR$ | $MeTMR$ |
| --- | --- | --- | --- | --- | --- | --- | --- | --- | --- |
| Peru | Jepelacio (High - 1123m) | 31 | 3.10 | 0.90 | 11 | 2.09 | 0.87 | 2.82 | 3 |
|  | Moyobamba (High - 955m) | 35 | 3.10 | 0.87 | 10 | 1.98 | 0.86 | 3.5 | 4 |
|  | Rio shilcayo basin (Mid - 444m) | 20 | 2.46 | 0.82 | 9 | 1.83 | 0.83 | 2.22 | 2 |
|  | Chazuta (Mid - 355m) | 23 | 2.53 | 0.81 | 9 | 1.59 | 0.72 | 2.56 | 2 |
|  | Pongo-Barrenquita road (Low - 191m) | 30 | 2.69 | 0.79 | 11 | 1.92 | 0.80 | 2.73 | 3 |
|  | Shucushyacu (Low - 165m) | 31 | 2.85 | 0.83 | 12 | 2.14 | 0.86 | 2.58 | 3 |
| French Guiana | Mont Itoupé (High - 826m) | 19 | 2.53 | 0.86 | 7 | 1.70 | 0.87 | 2.71 | 2 |
|  | Mont Galbao (High - 730m) | 19 | 2.31 | 0.79 | 7 | 1.67 | 0.86 | 2.71 | 2 |
|  | Tortue Mountain (Mid - 483m) | 15 | 2.54 | 0.94 | 8 | 1.79 | 0.86 | 1.88 | 2 |
|  | Kaw Mountain (Mid - 284m) | 12 | 2.03 | 0.82 | 6 | 1.42 | 0.79 | 2 | 1.5 |
|  | Nouragues station (Low - 54m) | 8 | 1.58 | 0.70 | 4 | 0.57 | 0.42 | 2 | 1 |
|  | Saut-Maripa (Low - 30m) | 5 | 0.12 | 0.76 | 3 | 0.78 | 0.71 | 1.67 | 1 |

In both regions, communities at high altitude exhibited a significantly higher overall phylogenetic clustering (NRI) than those at both low and mid-altitudes (p < 0.05), but no significant relationship between the more recent lineage-level patterns (NTI) and altitude class were found (all p > 0.2; see Supplementary Information, Table S3 and Fig. S6). PERMANOVA showed that altitude was a key driver of phylogenetic β-diversity, explaining 18% of the total variance (p = 0.03), with the strongest differences observed at low altitudes (low–mid: R² = 0.15, p = 0.04; low–high: R² = 0.17, p = 0.03; mid–high: p > 0.05). These results support the existence of phylogenetic differentiation between low and high altitudes, regardless of region. In Peru, this pattern was primarily driven by phylogenetic turnover, with nestedness contributing significantly less to overall phylogenetic β-diversity (β = –1.56, p < 0.001), whereas in French Guiana both turnover and nestedness contributed similarly to community dissimilarity (β = 0.09, p > 0.05; Fig. S7).

### The role of altitude in habitat filtering

#### Climatic variation and elevation: both temperature and precipitation decline with altitude

In both regions, daily temperature during sampling, annual mean temperature and annual precipitation decreased with altitude (p < 0.05), whereas no significant relationship was found for precipitation during sampling.

#### Ithomiini assemblages are not shaped by seasonality but remain structured by altitude

For the subset of sites in French Guiana sampled in both the rain and dry seasons, altitude class remained a significant factor influencing 59% of the taxa community variance (p = 0.007), whereas season (R² = 0.05, p = 0.8) and the interaction between season and altitude class (R² = 0.11, p = 0.9) were not significant. Taxa richness increased during the rain season compared to the dry season in low-altitude sites, but not mid and high-altitude sites (Fig. S8).

#### Predation pressure across elevation: no altitudinal pattern but a potential driver of diversity and mimicry-based mutualism

Predation rates on artificial butterfly models were relatively low across all sites (<0.06), but were consistent with values reported in previous studies (Chouteau et al., 2016). Although low absolute attack rates may limit inference on absolute predation intensity, the use of a large number of models per site provided a robust estimation of relative differences in predation pressure among sites and regions. Altitude had no significant effect on predation rates (p > 0.05), regardless of region (Fig. S9), although predation rates were generally higher in Peru. By contrast, predation rate had a significant positive effect on taxa richness (β = 20.68, p = 0.003; Fig. 2a), mimicry-ring richness (β = 14.96, p = 0.002; Fig. 2b), and the median number of taxa per mimicry ring (β = 18.13, p = 0.001; Fig. 2c), with a similar trend for the average number of taxa per mimicry ring (β = 6.16, p = 0.062). These effects were consistent across regions (see Supplementary Information).

**Figure 2.**
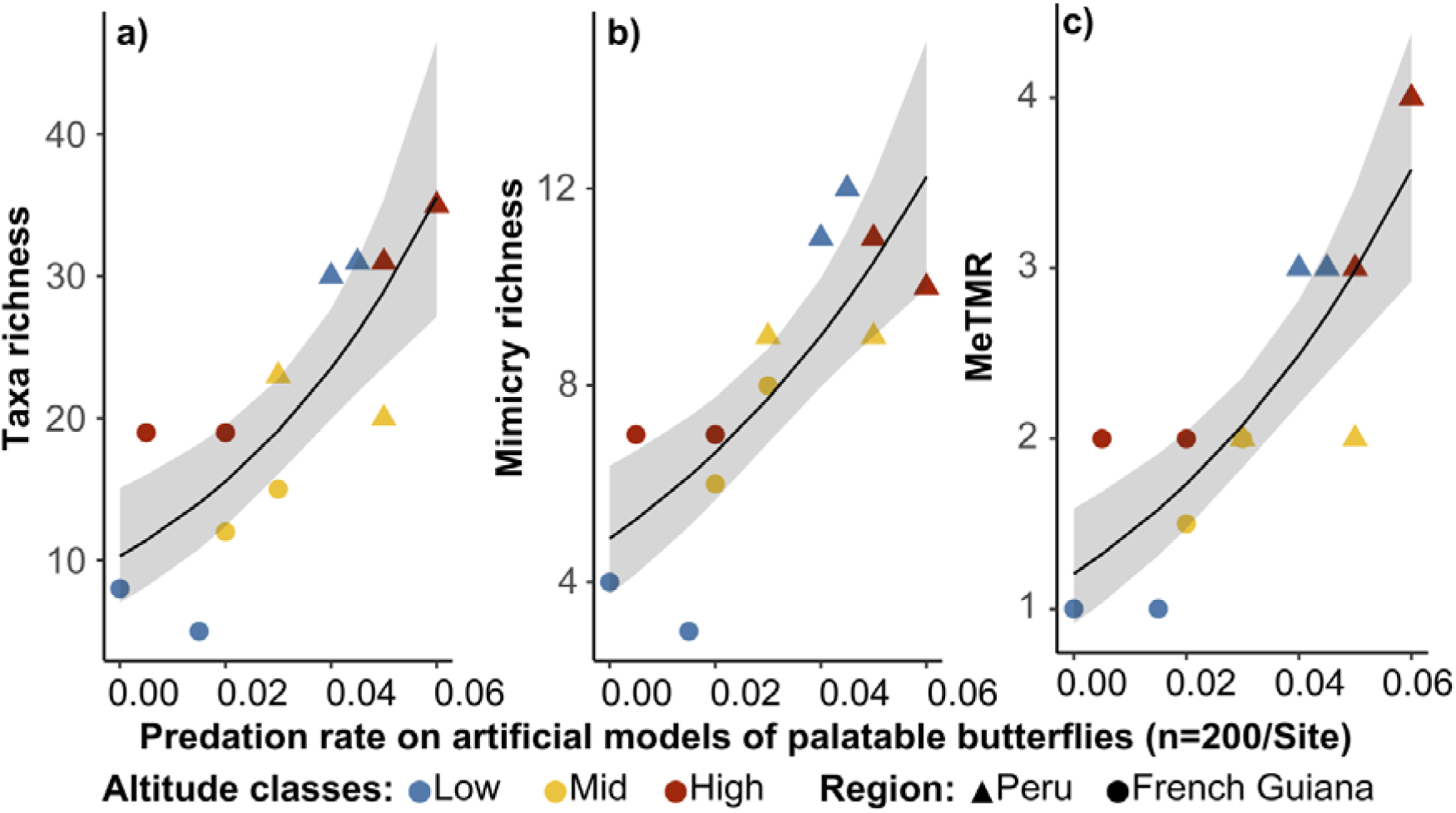
Species richness (a), mimicry ring richness (b) and the median number of taxa per mimicry ring (c) as a function of predation rate on artificial models of palatable butterflies (n = 200 per site). Colours indicate altitude classes and shape indicate regions. Lines represent the predictions from mixed generalized linear models showing significant effects (p < 0.05). Higher predation pressure was associated with increased diversity and stronger mimetic convergence.

#### Diversity begets diversity: potential host plant richness increases with elevation and seems to promote Ithomiini diversity

Each of the two stacked species distribution models (SSDMs) for Ithomiini butterflies and their potential host plants (Solanaceae and Gesneriaceae) in French Guiana demonstrated high AUC values, ranging from 0.75 to 0.92 (Table S4). Both models predicted similar patterns of species richness (Figs. 3a & 3b) following altitudinal gradients (Fig. 3c), with the exception that potential host plants exhibited higher richness along the coast. In both SSDMs, the maximum temperature of the warmest month was the most influential variable, explaining 47% for Ithomiini and 36% for potential host plants. The contributions of other variables were comparable, ranging between 12% and 21%. Richness of potential host plant and Ithomiini species were strongly correlated with altitude (R² = 0.87, p < 0.001; Fig. 4), both showing differences between altitudinal classes (see Fig. S10).

**Figure 3.**
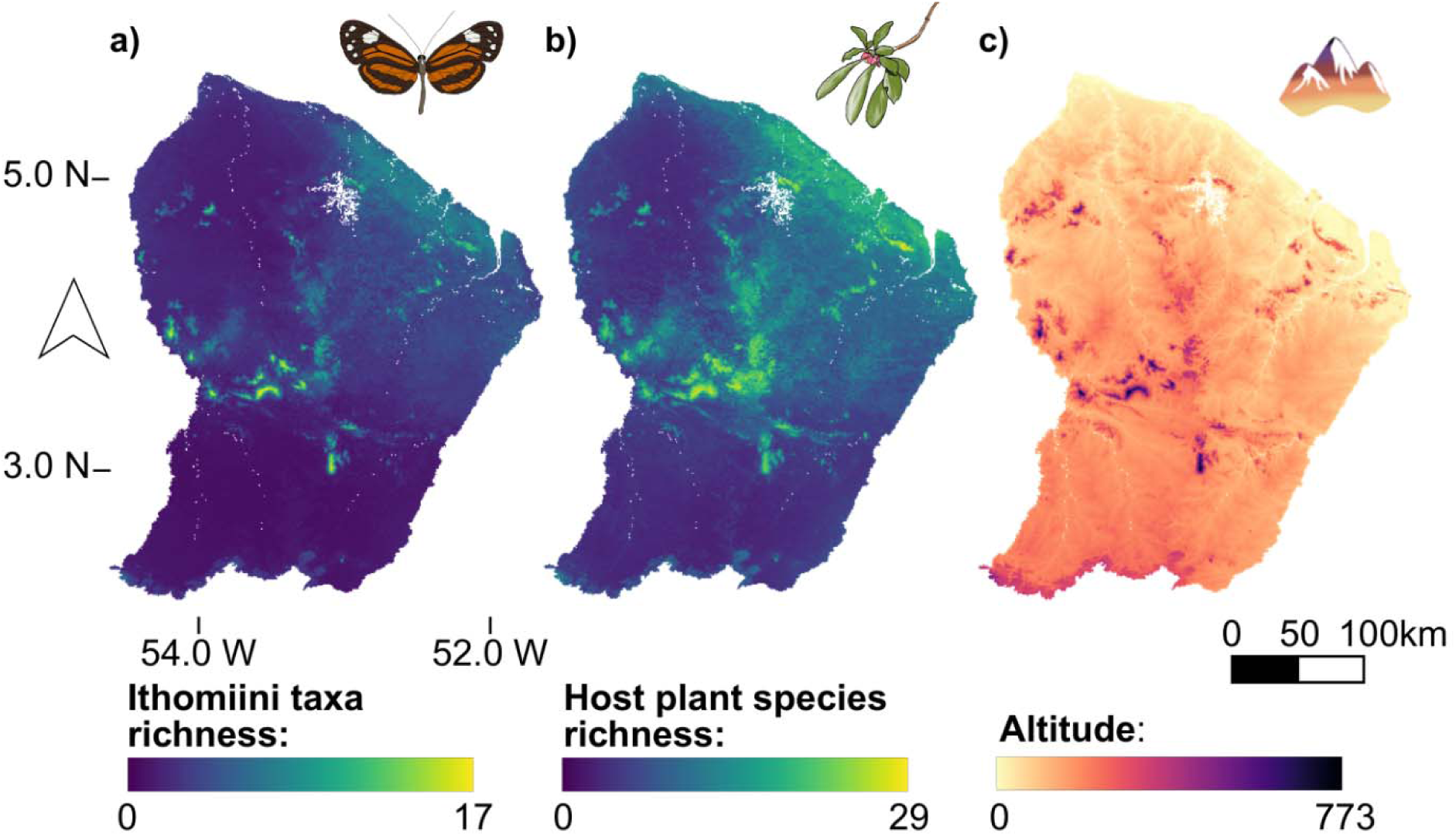
Stacked species distribution models of taxa richness for both (a) Ithomiini, and (b) their host plants (Solanaceae and Gesneriaceae) in French Guiana. (c) Elevation map of the region taken from WorldClim (Fick & Hijmans 2017). Spatial patterns of Ithomiini richness closely mirror those of their potential host plants, and both are correlated with the elevational structure of the landscape.

**Figure 4.**
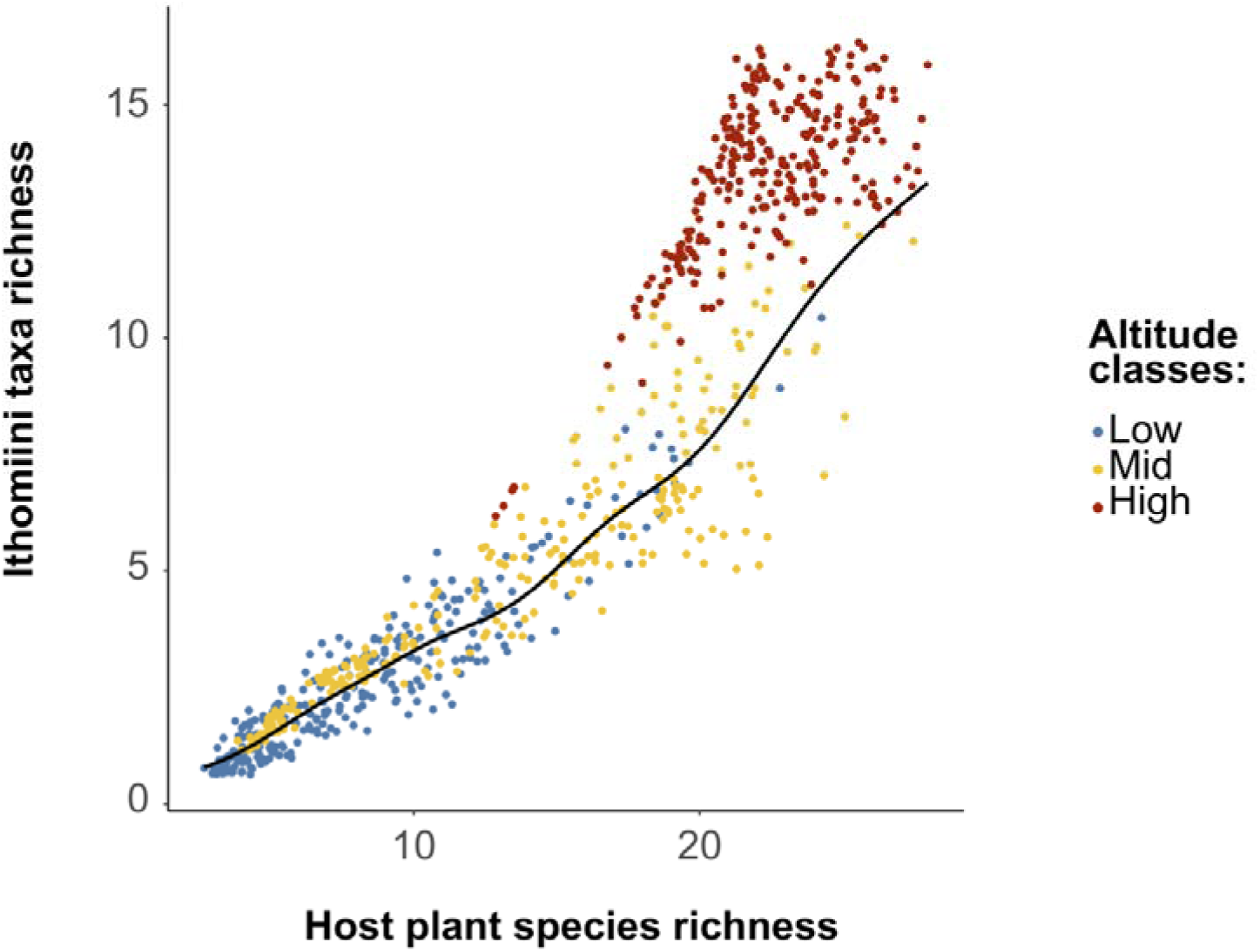
Relationship between Ithomiini species richness and host plant species richness in French Guiana, modelled using the SSDM R package. The generalized additive model (GAM) with a quasi-Poisson distribution was fitted on a dataset of N = 343 957 observations, and 277 randomly sampled points per altitude class (indicated by colours) are shown above. The black line represents the fitted generalized additive model across all observations (p < 0.05). Greater ecological opportunities, reflected by higher potential host plant richness, are associated with increased Ithomiini diversity along the elevational gradient.

#### Ecological factors shape Ithomiini assemblages across regions with differing evolutionary time

The combination of annual mean temperature, annual precipitation, predation rate and mean forest cover had a significant effect on Ithomiini communities independently of regional effects (Statistic = 8.17, p = 0.006). However, predation rate was the most significant (Statistic = 3.63, p = 0.005), with annual mean temperature having a near significant effect (Statistic = 2.18, p = 0.051) and annual precipitation and forest cover having no detectable effect on community dissimilarity (Statistic < 1.5, p > 0.05; see Supplementary Information).

## Discussion

Understanding the ecological mechanisms underlying the origin and maintenance of biodiversity has been a central focus of ecological and evolutionary research for decades, particularly in tropical regions where complex biotic interactions prevail. Biodiversity patterns observed along elevational gradients emerge from the interplay between processes operating at different spatial and temporal scales, notably the composition of regional species pool shaped by evolutionary history and dispersal, and local community assembly driven by environmental filtering and biotic interactions. By examining variation in ecological factors along elevational gradients and their effects on community structure, functional traits, and phylogenetic composition across distinct regions, this study provides insights into how regional and local processes jointly structure tropical biodiversity, and whether similar elevational patterns can emerge despite contrasting evolutionary histories.

Taxa and mimicry-ring richness were consistently higher in Peru than in French Guiana, suggesting that long-term historical and neutral processes, such as the Andean origin of Ithomiini and the extended time for diversification, have played a major role in the accumulation of both species and mimicry rings (Chazot et al., 2019). This regional difference highlights how evolutionary time and lineage persistence could shape community-level diversity. However, despite these marked differences in regional richness and evolutionary history, elevational patterns of richness were strikingly similar between regions, with both taxa richness (primarily at the subspecies level) and mimicry-ring richness increasing steadily from sea level to approximately 1300 m. This consistency suggests that similar local assembly processes operate along elevational gradients in both regions, filtering species from distinct regional pools in comparable ways, and this pattern supports the idea that, even with environmental filtering leading to phylogenetic clustering, high-altitude environments can promote diversification within clades pre-adapted to local conditions. Similar patterns have been reported in Andean Ithomiini communities (Chazot et al., 2014) and in other Neotropical butterflies such as *Heliconius* (Rosser et al., 2012), suggesting that tropical montane habitats may promote diversification rather than simply constrain diversity.

Community dissimilarity along elevational gradients was closely linked to the phylogenetic history of the tribe and was predominantly driven by phylogenetic turnover rather than nestedness. This suggests that environmental gradients filter different subsets of the regional species pool at different elevations, favoring lineage replacement rather than simple loss of richness. This is similar to findings from studies in other mountainous taxa, such as rodents in the Andes (Maestri & Patterson, 2016) and plant-pollinator networks in Greece (Minachilis et al., 2023), where ecological specialization and niche differentiation are key drivers of elevational species replacement. The stronger phylogenetic turnover observed in Peru likely reflects the combination of steeper environmental gradients and a larger, more phylogenetically diverse regional pool, shaped by long-term diversification in the Andes (Chazot et al., 2014; De-Silva et al., 2016). Additionally, the Andes constitute the origin of the Ithomiini (Chazot et al., 2019) and several montane lineages (e.g., *Hyalyris*, *Ithomia*, and the makrena clade of *Oleria*; De-Silva et al., 2016) are largely absent from French Guiana. In contrast, the assemblages recorded in French Guiana likely reflect a subset of lowland lineages that have secondarily colonized higher elevations, following more recent dispersal from western Amazonia (Chazot et al., 2019). The region’s more moderate topography and relatively continuous forest cover (Guitet et al., 2013) may have further limited opportunities for allopatric speciation, leading to weaker turnover and a comparatively higher nestedness. Despite these differences in species pool composition, the emergence of comparable elevational structuring in both regions indicates that similar ecological filters act repeatedly on distinct evolutionary lineages.

As widely reported in tropical systems, temperature and precipitation are key environmental drivers of biodiversity along both latitudinal and altitudinal gradients (e.g. Francis & Currie, 2003; Montejo-Kovacevich et al., 2020). In agreement with our second hypothesis, temperature emerged as a key abiotic variable associated with elevational changes in Ithomiini richness and community composition, although its effect in multivariate analyses was marginal. This suggests that temperature may primarily act as a broad-scale environmental constraint, shaping the pool of species able to persist along elevation gradients, rather than as a direct driver of local community differentiation (Sunday et al., 2012). This is supported by the observed increase in Ithomiini richness during the rain season in the lowland of French Guiana, when temperatures drop and precipitation increases, as well as by the marginal association between annual temperature and community composition in both regions. In contrast, forest cover and annual precipitation exhibited non-significant effects on communities, suggesting that their influence may be more indirect or dependent on specific ecological contexts such as habitat structure (Forister et al., 2010). Both temperature and precipitation can shape butterfly communities indirectly by influencing host plant richness and distribution (Kreft & Jetz, 2007; Tang et al., 2014), thereby modulating the ecological opportunities available for phytophagous specialists such as Ithomiini.

Consistent with the “diversity begets diversity” hypothesis, we found a positive correlation between potential host plant diversity and Ithomiini richness. This suggests that increased plant diversity could expand ecological opportunities for phytophagous specialists like Ithomiini, facilitating coexistence through niche partitioning and potentially promoting diversification (Janz et al., 2006; Lewinsohn & Roslin, 2008; Nylin & Janz, 2009). However, previous phylogenetic studies have shown limited evidence for phylogenetic tracking between Ithomiini and their Solanaceae hosts, suggesting that diversification may have resulted instead from repeated colonizations of chemically or ecologically similar host plant clades, followed by ecological specialization (Willmott & Freitas, 2006). In this regard, host plant shifts may have acted as key innovations triggering adaptive radiations (McClure & Elias, 2016). Nonetheless, it cannot be excluded that congruent richness patterns in Ithomiini and their potential host plants partly reflect shared responses to underlying environmental gradients.

Our findings also underscored the complex role of predation in shaping biodiversity patterns across both regions, supporting our third hypothesis. Notably, unlike other ecological factors examined, predation was not significantly correlated with altitude. Also, contrary to expectations that strong frequency-dependent selection would limit mimicry diversity (Chouteau et al., 2016), our results indicated that predation is positively correlated with taxa and mimicry-ring richness in both regions. Moreover, predation pressure was the most important factor explaining community differences, suggesting that biotic processes could more directly influence local community composition. This is consistent with evidence that generalist predators can support multiple mimetic morphs within diverse communities, as observed in *Heliconius* butterflies (Merrill et al., 2015) and suggests that predation may act as a driver of diversification, possibly through niche partitioning (Willmott et al., 2017). In tropical forests, predator communities are structured with different functional guilds exploiting distinct forest strata (Marra & Remsen, 1997; Walther, 2002). Such spatial heterogeneity in predation pressure can favor the persistence of multiple prey strategies by promoting segregation along microhabitat gradients. Vertical and microhabitat partitioning is well documented in Ithomiini butterflies (Beccaloni, 1997; DeVries et al., 1999; Hill, 2010; Willmott et al., 2017) where mimicry rings are not randomly distributed but tend to be associated with specific microhabitats. Spatial variation in predation pressure may reduce direct competition among co-occurring taxa and facilitate diversification, consistent with theoretical models showing that mimicry can promote species richness through predator-mediated spatial segregation (Aubier et al., 2017), thereby reconciling the paradox of high mimetic diversity under strong frequency-dependent selection. However, this pattern could also reflect the generally higher levels of diversity and abundance in both predators and butterflies, possibly due to broader environmental drivers typical of tropical diversity hotspots. Furthermore, strong predation pressure seems to promote mimicry-mediated mutualism by increasing the median of taxa richness per mimicry ring, supporting the notion that predator learning can enhance the stability and diversity of mimicry communities (Mallet & Gilbert, 1995; Willmott et al., 2017; Doré et al., 2023; Pérochon et al., 2025).

## Conclusion

In conclusion, we found higher taxa and mimicry richness in Peru, consistent with an Andean origin and longer evolutionary time. Despite these marked regional differences, elevational patterns of community structure were strikingly similar, with increases in richness and phylogenetic clustering at higher elevations, reflecting comparable habitat filtering along altitudinal gradients.

In addition to the potential influence of abiotic factors, particularly temperature, elevational variation in butterfly diversity was further associated with local biotic factors, including potential host plant richness and predation pressure. The positive relationship between host plant and butterfly richness is consistent with *diversity begets diversity*, whereby increased ecological opportunities promote species coexistence and diversification. In contrast to classical expectations that predation constrains diversity through strong stabilizing selection, our results suggest that predation pressure may instead act as a driver of diversification, shaping the fine scale structure of species and mimicry communities.

Despite strong differences in evolutionary history and regional species pools, Ithomiini communities exhibit remarkably consistent elevational patterns of richness, phylogenetic structure, and mimicry organization, suggesting that similar local ecological filtering repeatedly shape tropical butterfly communities across Neotropical montane systems.

## Acknowledgements

This work was funded by an “Investissement d’avenir” grant managed by the Center for the study of biodiversity in Amazonia (CEBA: ANR-10-LABX-25-01) to MM, by a French National Agency for Research grant (ANR SEXBOMB: ANR-21-CE02-0008) to MM, and benefited from a grant part of the France 2030 program and supported by the University of French Guiana and the Ministry of higher education and research (AIBSI: ANR-22-EXES-0005). We thank the Peruvian authorities for research permits (373-2017-SERFOR-DGGSPFFS) and the Gobierno Regional San Martín (PEHCBM permit: 012-2023/GRSM-PEHCBM-DMA-ACR-CE), and the French authorities for research permits in the Nouragues National Nature Reserve (no R03-2022-10-25-00004) and the Parc Amazonien de Guyane (PAG: no 1199-22). We would also like to thank URKU, INIBICO, Asociación de Protección de Fauna y Flora Alto Shilcayo, as well as Corina Amasifuen, Genaro Montilla and Flor Montilla Amasifuen, for all their help in Peru. In French Guiana, we would like to thank the CNRS Nouragues research field station (funded by AnaEE ANR-11-INBS-0001 and Labex CEBA ANR-10-LABX-25-01), the OHM Oyapock (funded by Labex DRIIHM/IRDHEI ANR-11-LABX-0010), and the late Denis Armagnac (Camp Bonaventure, French Guiana) for their assistance.

## Author Contributions

MM designed the study and acquired funding. RF & AR did the fieldwork, with the help of CP, GCP, RMP, LPC, NP, OS, SG, ME & MM. SG and MM secured permits in Peru and French Guiana. RF analyzed the data, and RF, ME & MM wrote the paper with contributions from all authors. All authors read and approved the final manuscript.

## Statement on inclusion

Our study brings together authors from a number of different regions/countries, including researchers based where the study was carried out. These diverse authors contributed in the early stages of the project, ensuring that diverse perspectives were considered.

## Data and code availability

Data will be made available on Zenodo before publication. https://doi.org/10.5281/zenodo.18302248

