## Supplementary information for "Repeated trends in altitudinal gradients of diversity: how habitat filtering and biotic interactions structure ecological communities"

[*]Co-last authors

*French abstract*

Comprendre comment la biodiversité est structurée le long des gradients altitudinaux tropicaux nécessite de dissocier le rôle de l'histoire évolutive régionale de celui des processus écologiques opérant à l'échelle locale. Dans cette étude, les communautés de papillons Ithomiini ont été étudiées le long de gradients altitudinaux répétés dans deux régions néotropicales aux trajectoires évolutives contrastées — les Andes amazoniennes et le bouclier guyanais — afin de déterminer si des schémas altitudinaux similaires de structure taxonomique, mimétique et phylogénétique émergent malgré des pools d'espèces régionaux distincts, et si des facteurs abiotiques et biotiques contribuent à façonner ces schémas.

Malgré des différences marquées de richesse régionale, des patrons altitudinaux remarquablement similaires émergent dans les deux régions. La richesse taxonomique et mimétique augmente avec l'altitude et s'accompagne d'un regroupement phylogénétique plus prononcé, indiquant que des processus de filtrage de l'habitat comparables semblent opérer le long des gradients altitudinaux, indépendamment du contexte régional. La β-diversité phylogénétique semble principalement déterminée par le turnover des lignées, en particulier dans les Andes, soulignant le rôle des gradients altitudinaux dans le remplacement de lignées phylogénétiquement distinctes. Ces résultats suggèrent que l'altitude exerce un effet fort et répétable sur la structure des communautés, avec un filtrage environnemental pouvant agir localement sur des pools d'espèces régionaux différents.

Les facteurs abiotiques, tels que la température, semblent contraindre la distribution des espèces à grande échelle spatiale, tandis que les interactions biotiques semblent principalement opérer à l'échelle locale. En particulier, la diversité des papillons est positivement associée à la richesse potentielle en plantes-hôtes et à la pression de prédation, indiquant que les interactions écologiques contribuent à façonner la composition des communautés locales une fois les contraintes environnementales générales prises en compte. En intégrant structure phylogénétique, interactions biotiques et gradients environnementaux dans des régions aux histoires évolutives contrastées, cette étude met en évidence la manière dont les pools régionaux et le filtrage écologique local interagissent pour structurer la biodiversité tropicale, et souligne que des processus d'assemblages altitudinaux similaires pourraient survenir indépendamment le long des gradients altitudinaux néotropicaux.

*Spanish abstract*

Comprender cómo se estructura la biodiversidad a lo largo de los gradientes altitudinales tropicales requiere disentanglar el papel de la historia evolutiva regional y el de los procesos ecológicos que operan a escala local. En este estudio, se analizaron comunidades de mariposas Ithomiini a lo largo de gradientes altitudinales repetidos en dos regiones neotropicales con trayectorias evolutivas contrastantes — los Andes amazónicos y el Escudo Guayanés — para evaluar si emergen patrones altitudinales similares de estructura taxonómica, mimética y filogenética a pesar de contar con pools regionales de especies distintos, y si factores abióticos y bióticos contribuyen a estructurar estos patrones.

A pesar de marcadas diferencias en la riqueza regional, emergen patrones altitudinales notablemente similares en ambas regiones. La riqueza taxonómica y mimética aumenta con la altitud y se acompaña de un mayor agrupamiento filogenético, lo que indica que procesos comparables de filtrado del hábitat parecen operar a lo largo de los gradientes altitudinales, independientemente del contexto regional. La β-diversidad filogenética parece estar determinada principalmente por el recambio de linajes, en particular en los Andes, lo que resalta el papel de los gradientes altitudinales en la sustitución de linajes filogenéticamente distintos. Estos resultados sugieren que la altitud ejerce un efecto fuerte y repetible sobre la estructura de las comunidades, con el filtrado ambiental a escala local actúa potencialmente sobre distintos pools de especies regionales.

Los factores abióticos, como la temperatura, limitan la distribución de las especies a escalas espaciales amplias, mientras que las interacciones bióticas parecen operar principalmente a escala local. En particular, la diversidad de mariposas se asocia positivamente con la riqueza potencial de plantas hospedadoras y con la presión de depredación, lo que indica que las interacciones ecológicas contribuyen a modelar la composición de las comunidades locales una vez consideradas las restricciones ambientales generales. Al integrar la estructura filogenética, las interacciones bióticas y los gradientes ambientales en regiones con historias evolutivas contrastantes, este estudio pone de relieve cómo los pools regionales de especies y el filtrado ecológico local interactúan para estructurar la biodiversidad tropical, y subraya que procesos de ensamblaje altitudinal similares podrían ocurrir independientemente a lo largo de los gradientes altitudinales neotropicales.

### Methods

#### *Study sites*

In Peru, the sites consisted of Shucushyacu (5°59’40”S, 75°52’18”W, altitude (alt.) 165 m), Pongo de Caynarachi - Barranquita road (Carratera Pelejo Papaplaya, 6°17’22”S, 76°12’44”W, alt. 191 m, referred to as Pongo-Barranquita), Chazuta (6°32’39”S, 76°07’06”W, alt. 355 m), the Rio Shilcayo basin near Tarapoto (6°27’47”S, 75°52’18”W, alt. 444 m, referred to as Rio Shilcayo), Moyobamba (6°04’22”S, 76°58’15”W, alt. 955 m), and Jepelacio (6°05’30”S, 76°58’05”W, alt.

1123 m). In French Guiana, sites consisted of the Nouragues Nature Reserve (CNRS station Pararé 4°02'17"N, 52°40'22"W, alt. 54 m, referred to as Nouragues), Saut-Maripa (3°52'58"N, 51°51'53"W, alt. 30 m), Kaw Mountain (4°33'26"N, 52°08'54"W, alt. 284 m), Grande Tortue Mountain (4°17'50"N, 52°21'34"W, alt. 483 m, referred to as Tortue mountain), Mont Galbao (3°36'06"N, 53°16'21"W, alt. 730 m), and Mont Itoupé (3°01'22"N, 53°05'18"W, alt. 826 m). Sampling was done between June and September 2022 in Peru, and between October and December 2022 and November 2023 in French Guiana. Three sites in French Guiana (one from each altitude) were re-sampled during the rain season of 2024 (March-April) to assess the potential influence of seasonality on community composition. This additional data was used exclusively to test seasonal effects and were not included in any other analyses.

Before conducting any statistical tests, rarefaction curves were used to assess whether sampling effort was sufficient across altitudinal ranges (iNEXT.3D R package; Chao et al., 2021). Since sample coverage reached an asymptote, it can be assumed that sampling was adequate for further analyses (Fig. S1).

*Spatial patterns of diversity and phylogenetic community structure in Ithomiini*

To test whether species and mimicry-ring richness differed between French Guiana and Peru, and whether altitudinal patterns in diversity were consistent across regions, generalized linear mixed models (GLMMs) with Poisson distributions were applied separately for species richness and mimicry-ring richness. For each response variable, two models were fitted: the first included region as a fixed effect and altitude class as a random effect, to assess regional differences while accounting for altitudinal variability; the second included altitude class as a fixed effect and region as a random effect, to evaluate altitudinal patterns across regions.

To test whether Ithomiini communities exhibit similar altitudinal patterns of species and mimicry-ring composition in both regions, permutational multivariate analysis of variance (PERMANOVA; R package vegan; Oksanen et al., 2024) was applied separately to taxa and mimicry-rings Bray–Curtis dissimilarity matrices between sites. For each matrix, the model included altitude class, region, and their interaction as explanatory variables, in order to assess whether variation in community structure was primarily driven by altitude, by region, or by region-specific altitudinal effects, and post-hoc pairwise comparisons were done using the pairwise Adonis R package (Martinez Arbizu, 2020). Because PERMANOVA assumes homogeneity of multivariate dispersion among groups, this assumption was assessed. For both taxa and mimicry-ring community matrices, no significant differences in multivariate dispersion were detected among altitude classes (*T*: F = 0.56, p = 0.78; *MR*: F = 4.22, p = 0.053) or between regions (*T*: F = 0.23, p = 0.63; *MR*: F = 0.59, p = 0.44), indicating that PERMANOVA results for these main effects reflect differences in community composition rather than differences in within-group variability. For both taxa and mimicry-ring matrices, the Altitude × Region interaction showed a significant dispersion test (p = 0.001). However, this result was associated with near-zero residual variance reflecting the limited number of sites per interaction level rather than biologically meaningful differences in dispersion. Consequently, this interaction was not interpreted, and PERMANOVA results were primarily evaluated with respect to the main effects of altitude and region.

Phylogenetic α- and β-diversity indices were compared using the most recent phylogeny of the Ithomiini tribe (Chazot et al., 2019). Faith's phylogenetic diversity (PD) was first used to quantify the total branch length across observed species in each community. Because Faith's PD is not statistically independent from species richness, two additional indices—net relatedness index (NRI) and nearest taxon index (NTI)—were used to evaluate patterns of phylogenetic clustering (higher values) and dispersion (lower values).

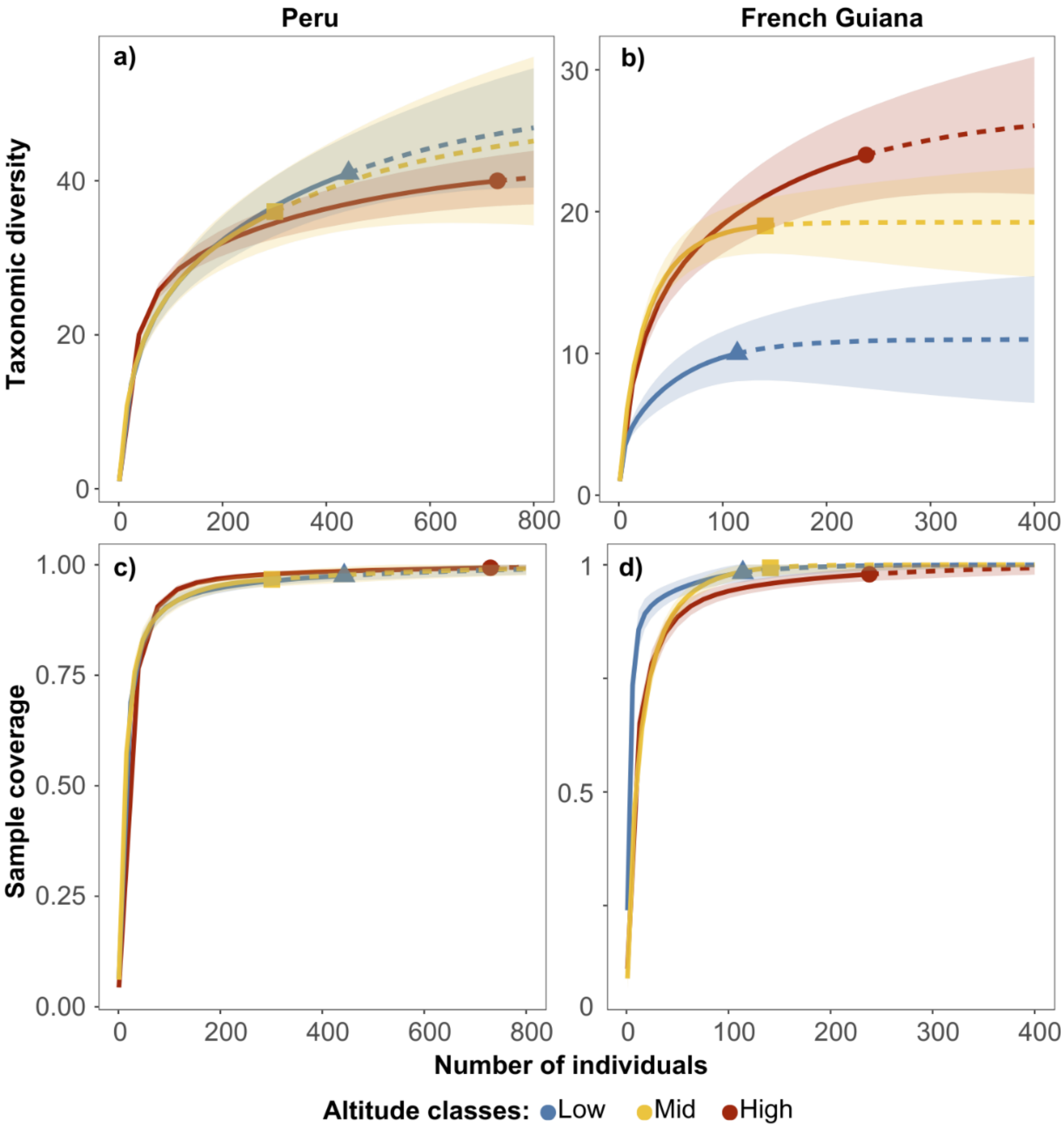


**Figure S1.** Rarefaction curves showing taxa richness (a, b) and sample coverage (c, d) as a function of abundance, coloured by altitude classes. Panels (a) and (c) correspond to Peru, while panels (b) and (d) correspond to French Guiana. Sample coverage reaches an asymptote across altitude classes and regions, indicating that sampling effort was sufficient to reliably characterize community richness.

NRI was calculated based on the standardized effect size mean (pairwise) phylogenetic distances (SES-MPD), which reflects deep phylogenetic structure, whereas NTI was calculated using the mean nearest neighbour phylogenetic distance (SES-MNTD), a metric sensitive to terminal phylogenetic patterns (Webb, 2000; Webb et al., 2002). Standardized effect sizes, for both MPD and MNTD, were calculated for a distribution of 10 000 null values, generated by randomly shuffling tip labels on the phylogeny with the R package picante (Kembel et al., 2010).

To evaluate whether patterns of phylogenetic clustering with elevation were consistent between the Peruvian Andes and French Guiana, as a result of similar ecological processes, linear mixed models (LMMs) were fitted separately for the NRI and the NTI. In both models, the region was included as a random effect to account for broad-scale biogeographic differences. Importantly, the models were weighted by the standard deviation of the null model, which ensured that estimates were adjusted for uncertainties inherent to the null model simulation. This approach also allowed the inclusion of both significant and non-significant site-level values relative to the neutral expectation (i.e. 0), thereby retaining the full range of variation in phylogenetic structure across sites.

For phylogenetic community analyses, homogeneity of multivariate dispersion was assessed based on UniFrac distance matrices. No significant differences in dispersion were detected among altitude classes ($F = 0.56$, $p = 0.58$) or between regions ($F = 0.59$, $p = 0.55$), indicating that PERMANOVA results for these main effects are not driven by heterogeneity in within-group dispersion. As observed for taxonomic and mimicry-ring community matrices, the Altitude × Region interaction showed a significant dispersion ($p = 0.001$). Consequently, this interaction term was not interpreted, and phylogenetic PERMANOVA results were primarily evaluated with respect to the main effects of altitude and region. The comparison of the relative contribution of phylogenetic turnover (i.e. lineage replacement between sites) and nestedness (i.e. differences due to the gain or loss of lineages, where one community is a subset of another) to community dissimilarity was assessed with a generalized linear model using a Tweedie distribution (R package cplm; Zhang, 2013). This distribution was selected for its suitability in modelling non-normal, strictly positive data with potential heteroscedasticity. The index parameter was estimated at $p = 1.41$, indicating a distributional shape intermediate between Poisson and gamma. Model assumptions were checked using diagnostic plots (residuals vs. fitted, histogram, QQ-plot) and by testing for heteroscedasticity via a Spearman correlation between absolute residuals and fitted values ($\rho = -0.16$, $p = 0.21$).

*The role of altitude in habitat filtering*

*Climatic variation and elevation: temperature and precipitation gradients*

Rainfall data for the sampling days were obtained from AgERA 5 (Boogaard et al., 2020) using the R package ag5Tools (Brown et al., 2023). To visualize changes in precipitation along the altitudinal gradient, generalized linear models (GLMs) were performed separately for both Peru and French Guiana after checking for overdispersion. In French Guiana, the Mont Galbao site was excluded due to its status as an outlier. This site exhibited an exceptionally high rainfall value (139.60 mm) compared to the other sites, where rainfall ranged from 1.45 mm to 19.12 mm.

*Effect of seasonality and short-term climatic conditions on Ithomiini assemblages*

A PERMANOVA was conducted to test whether taxa community composition was influenced by seasonality. The model included altitude class, season (dry vs. rain), and their interaction. No significant differences in dispersion were detected among altitude classes ($F = 0.80$, $p = 0.48$) or between seasons ($F = 0.87$, $p = 0.35$), indicating that PERMANOVA results for these main effects are not driven by heterogeneity in within-group dispersion. The altitude × season interaction showed a significant dispersion ($p = 0.001$). Consequently, this interaction term was not interpreted, and PERMANOVA results were primarily evaluated with respect to the main effects of altitude and season. To assess the influence of ecological factors (both abiotic and biotic) on Ithomiini species and mimicry-ring composition, preliminary analyses were conducted to ensure that short-term weather conditions during data collection did not bias our sampling. An initial mixed multivariate distance matrix regression (MDMR packages in R, McArtor et al., 2016) analysis was performed with region as a random factor to test whether climatic conditions at the time of sampling—specifically, average daytime temperature (06:00–19:00; measured with HOBO sensors) and rainfall (AgERA5)—influenced species community composition. Although the omnibus test was not significant, indicating no overall correlation between these short-term climatic variables and community variations, the average daytime temperature showed a significant effect likely reflecting a weak but detectable influence on community composition. However, a subsequent spearman correlation test revealed a significant relationship between average daytime temperature during sampling and annual temperature (Spearman correlation = 0.81, $p = 0.001$). This result suggests that the observed effect was not due to short-term climatic fluctuations but rather reflected broader climatic patterns.

*Predation pressure across elevation and its influence on diversity and mutualism*

Models were based on locally abundant palatable species (*Pierella hyceta* in Peru and *Pierella astyoche* in French Guiana; Fig. S2). The butterfly models were constructed using modelling clay (Van Aken) for the body, and high-resolution wing patterns printed on matte photographic paper (EPSON Double-Sided Matte Paper C13S041569) using a Canon imagePRESS C265 printer. To ensure colour accuracy, printed wing patterns were verified using colorimetric analysis with the X-Rite ColorChecker Classic Mini. The use of two palatable and ubiquitous butterfly species was intended to minimize major biases in the estimation of predation rates at each locality. First, selecting a cryptic palatable species eliminates biases associated with predator learning, which may vary across predator communities and locations (Endler & Mappes, 2004; Mappes et al., 2005). Second, the use of widespread species helps reduce potential effects of neophobia and dietary conservatism, which can lead to reduced attack rates on novel or rare prey (Marples et al., 1998; Marples & Kelly, 1999; Szabo & Ringler, 2022). This approach follows established protocols used in similar field experiments aiming to assess visual predation, where only beak-shaped marks or clear tearing attributable to bird attacks are considered valid predation events, excluding ambiguous or arthropod-related damage (Arias et al., 2016; Chouteau et al., 2016).

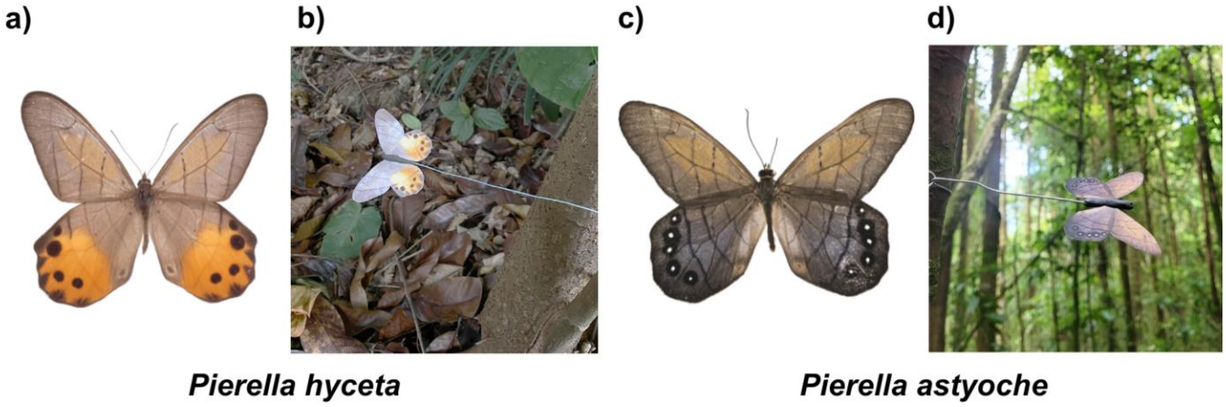


**Figure S2.** Photographs of (a, b) *Pierella hyceta* from Peru and (c, d) *Pierella astyoche* from

French Guiana. Panels (a) and (c) show butterflies on a white background, while panels (b) and (d) display models in situ for predation tests. Palatable and locally abundant butterfly species were used as experimental models for estimating background avian predation pressure under natural conditions.

Models were placed every 10 meters along transects at varying heights (at approximately 50 cm and 2 m) to mimic Ithomiini flight height and left in situ for 3 days (Arias et al., 2016; Chouteau et al., 2016) in the same sites where butterflies were collected. Although researcher presence could have reduced predator activity in the immediate vicinity, this disturbance was minimal, spatially limited, and consistent across all sites, and therefore unlikely to bias comparative estimates of predation pressure. After retrieval, predation marks by avian predators were recorded by the same two observers to avoid bias (see Fig. S3 for examples).

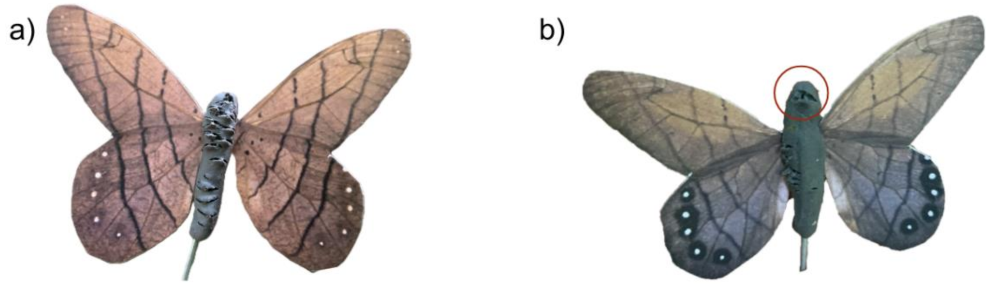


**Figure S3.** Examples of predation marks on experimental models: (a) marks left by insects; (b) a bird beak mark (circled in red). Only avian predation was considered, and bird attack marks were recorded by the same two observers to avoid observer bias.

*Host plant diversity across elevation: the "diversity begets diversity" hypothesis*

Climatic variables were selected through a two-step process to minimize multicollinearity. First, a Pearson correlation matrix was used to identify and exclude highly correlated variables (threshold $r > 0.7$). This was followed by a variance inflation factor (VIF) analysis, applying a cut-off value below 10, as recommended by Guisan et al. (2017). The final set of retained variables included:

(1) Bio1 – annual mean temperature; (2) Bio2 – mean diurnal temperature range; (3) Bio4 – temperature seasonality, quantified as the standard deviation of monthly temperature values multiplied by 100; (4) Bio18 – precipitation of the warmest quarter; and (5) Bio19 – precipitation of the coldest quarter.

To reduce potential biases associated with heterogeneous sampling effort, the selection of pseudo-absence points was refined using the target-group background approach (Phillips et al., 2009; Barber et al., 2022). This approach constructs a background layer that mirrors actual sampling intensity by generating a density surface based on occurrences from taxa surveyed using comparable methods within the same region. A two-dimensional kernel density estimation was performed following the procedure described by Barber et al. (2022). The resulting raster, matching the spatial extent and resolution (30 arc-seconds) of the environmental predictors, was normalized to a scale from 1 to 20 as recommended by Elith et al. (2010), thereby increasing the likelihood of selecting background points in more intensively sampled areas (Fig. S4). This procedure helps mitigate artifacts caused by spatial sampling bias in species distribution models.

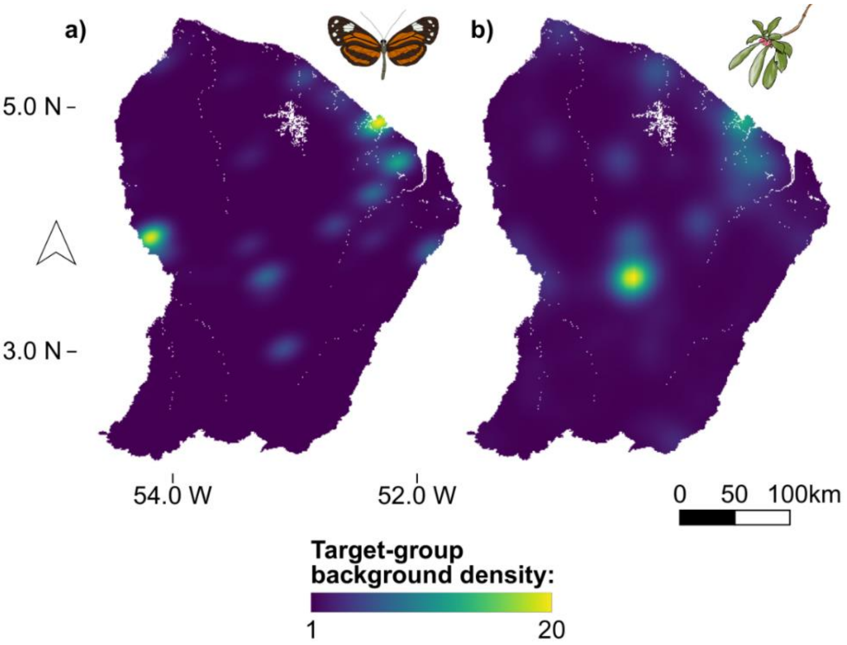


**Figure S4.** Kernel density maps of occurrence records used for the target-group background with (a) density of occurrences for Ithomiini, and (b) density of occurrences for their host plants (Solanaceae and Gesneriaceae). Density values are scaled between 1 and 20. Strong spatial heterogeneity in sampling effort supports the use of target-group background corrections in species distribution modelling.

The stacked species distribution model for Ithomiini butterflies was generated using the SSDM R package (Schmitt et al., 2017). All modelling algorithms available within the SSDM package were implemented, including artificial neural networks (ANN), classification tree analysis (CTA), generalized additive models (GAM), generalized boosted regression models (GBM), generalized linear models (GLM), multivariate adaptive regression splines (MARS), maximum entropy (MXT), random forest (RF), and support vector machines (SVM).. Species occurrences were filtered to include only taxa with a minimum of 10 occurrences, resulting in 21 Ithomiini species (Table S1) and 47 potential host plant species (Table S2) retained for modelling. A total of 10 replicates per species were run using 10-fold cross-validation. Ensemble predictions for each taxon were obtained by retaining algorithms that met the ensemble selection criterion (AUC $\geq$ 0.7; ensemble.thresh = 0.7) and, when applicable, combining retained algorithms using the SSDM ensemble procedure. Species richness was then estimated using probabilistic stacking (pSSDM), by summing continuous occurrence probabilities across taxa.

**Table S1.** Performance metrics (AUC and Kappa) of ensemble species distribution models and number of occurrence records (n) for each Ithomiini species.

| Species | AUC | Kappa | n |
|---|---|---|---|
| *Callithomia alexirrhoe* | 0.87 | 0.41 | 15 |
| *Callithomia lenea* | 0.88 | 0.41 | 21 |
| *Ceratinia cayana* | 0.87 | 0.46 | 13 |
| *Ceratinia neso* | 0.85 | 0.36 | 24 |
| *Hyposcada anchiala* | 0.91 | 0.53 | 32 |
| *Hyposcada dujardini* | 0.91 | 0.43 | 28 |
| *Hyposcada zarepha* | 0.88 | 0.36 | 23 |
| *Hypothyris euclea* | 0.85 | 0.38 | 12 |
| *Hypothyris ninonia* | 0.90 | 0.52 | 82 |
| *Hypothyris vallonia* | 0.90 | 0.49 | 10 |
| *Mechanitis polymnia* | 0.93 | 0.60 | 123 |
| *Melinaea ludovica* | 0.90 | 0.54 | 115 |
| *Melinaea mediatrix* | 0.83 | 0.32 | 35 |
| *Melinaea mneme* | 0.87 | 0.44 | 111 |
| *Methona grandior* | 0.84 | 0.33 | 33 |
| *Methona megisto* | 0.84 | 0.33 | 18 |
| *Napeogenes rhezia* | 0.85 | 0.35 | 23 |
| *Oleria aegle* | 0.92 | 0.53 | 52 |
| *Oleria astrea* | 0.84 | 0.35 | 16 |
| *Oleria flora* | 0.94 | 0.53 | 15 |
| *Oleria ilerdina* | 0.92 | 0.56 | 38 |

**Table S2.** Performance metrics (AUC and Kappa) of ensemble species distribution models and number of occurrence records (n) for each potential host plant species.

| **Species** | **AUC** | **Kappa** | **n** |
|---|---|---|---|
| *Brunfelsia guianensis* | 0.76 | 0.29 | 112 |
| *Brunfelsia martiana* | 0.88 | 0.47 | 10 |
| *Capsicum chinense* | 0.90 | 0.45 | 11 |
| *Capsicum frutescens* | 0.91 | 0.46 | 11 |
| *Columnea calotricha* | 0.78 | 0.29 | 87 |
| *Columnea oerstediana* | 0.88 | 0.45 | 113 |
| *Columnea sanguinea* | 0.87 | 0.39 | 66 |
| *Drymonia coccinea* | 0.77 | 0.32 | 231 |
| *Drymonia psilocalyx* | 0.92 | 0.54 | 62 |
| *Drymonia serrulata* | 0.75 | 0.29 | 109 |
| *Lycianthes pauciflora* | 0.87 | 0.39 | 56 |
| *Markea coccinea* | 0.74 | 0.25 | 98 |
| *Markea formicarum* | 0.84 | 0.32 | 16 |
| *Markea longiflora* | 0.85 | 0.34 | 30 |
| *Markea sessiliflora* | 0.80 | 0.25 | 29 |
| *Physalis angulata* | 0.84 | 0.35 | 51 |
| *Physalis pubescens* | 0.82 | 0.34 | 33 |
| *Solanum americanum* | 0.87 | 0.37 | 29 |
| *Solanum anceps* | 0.86 | 0.42 | 74 |
| *Solanum arboreum* | 0.85 | 0.37 | 17 |
| *Solanum asperum* | 0.84 | 0.38 | 84 |
| *Solanum coriaceum* | 0.84 | 0.39 | 71 |
| *Solanum costatum* | 0.90 | 0.46 | 14 |
| *Solanum crinitum* | 0.83 | 0.32 | 30 |
| *Solanum endopogon* | 0.86 | 0.38 | 61 |
| *Solanum fulvidum* | 0.86 | 0.39 | 24 |
| *Solanum jamaicense* | 0.87 | 0.40 | 39 |

| | | | |
|---|---|---|---|
| *Solanum leucocarpon* | 0.84 | 0.45 | 180 |
| *Solanum monachophyllum* | 0.84 | 0.32 | 13 |
| *Solanum morii* | 0.92 | 0.54 | 58 |
| *Solanum oppositifolium* | 0.82 | 0.31 | 36 |
| *Solanum paludosum* | 0.85 | 0.36 | 13 |
| *Solanum rubiginosum* | 0.84 | 0.35 | 53 |
| *Solanum rugosum* | 0.81 | 0.35 | 94 |
| *Solanum schlechtendalianum* | 0.82 | 0.36 | 77 |
| *Solanum schomburgkii* | 0.86 | 0.34 | 10 |
| *Solanum semotum* | 0.85 | 0.33 | 20 |
| *Solanum sendtnerianum* | 0.88 | 0.35 | 22 |
| *Solanum splendens* | 0.81 | 0.33 | 13 |
| *Solanum stramonifolium* | 0.83 | 0.37 | 70 |
| *Solanum subinerme* | 0.88 | 0.49 | 122 |
| *Solanum tegore* | 0.79 | 0.27 | 37 |
| *Solanum torvum* | 0.86 | 0.37 | 43 |
| *Solanum tricuspidatum* | 0.88 | 0.46 | 29 |
| *Solanum uncinellum* | 0.87 | 0.39 | 26 |
| *Solanum velutinum* | 0.81 | 0.33 | 74 |
| *Witheringia solanacea* | 0.85 | 0.39 | 16 |

To visualize whether both host plant and Ithomiini richness varied along elevational gradients, a linear discriminant analysis (LDA) was first conducted on 277 randomly sampled values (corresponding to the number of high-elevation pixels) to explore whether metrics of richness could visually discriminate between altitude classes. Due to violations of LDA assumptions—namely multivariate normality and homogeneity of covariance matrices—this analysis was used strictly for exploratory visualisation.

*Ecological predictors of Ithomiini assemblages*

Multicollinearity among variables was assessed using variance inflation factor (VIF) analysis, and variables with VIF values exceeding 10, such as altitude, were excluded from subsequent analyses. Forest cover was calculated using five photographs per site, captured with a Sony Alpha 7 III equipped with a wide-angle lens (Sony 18mm) and analysed in ImageJ (Schindelin et al., 2012) to calculate the proportion of pixels identified as "forest canopy". Mixed MDMR models were applied to assess the influence of ecological predictors on Ithomiini composition, with region included as a random effect (Anderson, 2001; Zapala & Schork, 2012; Oyesola et al., 2024). This multivariate approach models the variation in community dissimilarities (Bray-Curtis) as a function of all retained environmental variables simultaneously (annual mean temperature, annual precipitation, predation rate and mean forest cover), allowing the relative contribution of each variable to be evaluated.

### Results

*Spatial patterns of diversity and phylogenetic community structure in Ithomiini: the effect of evolutionary history does not override the influence of altitude*

Taxa richness ($S_T$) and mimicry richness ($S_{MR}$) were significantly higher in Peru than in French Guiana ($S_T$: Peru - $\beta = 15.8 \pm 2.62$, $z = 6.02$, $p < 0.001$, French Guiana (Intercept) - $\beta = 12.68 \pm 2.43$, $z = 5.21$, $p < 0.001$; $S_{MR}$: Peru - $\beta = 4.5 \pm 1.64$, $z = 2.74$, $p = 0.006$, French Guiana (Intercept) - $\beta = 5.83 \pm 0.99$, $z = 5.9$, $p < 0.001$). Results show an increase in taxa richness at higher altitudes, and this is significant between low and high altitude ($S_T$: low-high: $\beta = 9.64 \pm 3.11$, $z = 3.1$, $p = 0.005$), but not with mid-altitudes ($S_T$: low-mid: $\beta = 2.42 \pm 2.76$, $z = 0.88$, $p = 0.655$; mid-high: $\beta = 7.22 \pm 3.13$, $z = 2.31$, $p = 0.054$). A similar, though non-significant, trend was observed for mimicry richness between low and high altitude ($S_{MR}$: low-high: β = 1.81 ± 1.98, z = 0.91, p = 0.63). PERMANOVA showed that altitude was a key driver of taxa community structure, explaining 19% of the total variance ($R^2$ = 0.19, F = 2.24, df = 2,6, p = 0.016). A similar pattern was observed in mimetic communities, with altitude also accounting for 19% of the variance ($R^2$ = 0.19, F = 2.32, df = 2,6, p = 0.039). Pairwise comparisons revealed significant differences between high and low altitudes for both taxa (T) and mimicry ring (MR) communities (T: $R^2$ = 0.10, F = 7.25, df = 1,4, p = 0.037; MR: $R^2$ =0.20, F = 9.67, df = 1,4, p = 0.02), whereas differences with mid-altitudes were not significant (p > 0.05), except for differences in mimetic communities between low and mid-altitudes ($R^2$ = 0.19, F = 4.59, df = 1,4, p = 0.04). Species diversity, as measured by the Shannon index ($H$), showed similar patterns between Peru and French Guiana across altitude classes. In Peru, $H_T$ was highest at high-altitude sites (3.10) and decreased slightly at mid-altitude (2.46–2.53) and low-altitude sites (2.69–2.85). In French Guiana, $H_T$ remained relatively stable between high and mid-altitude sites (2.03–2.54), but showed a sharp decline in low-altitude sites (0.12–1.58). For mimicry rings ($H_{MR}$), values remained relatively stable across altitudes in Peru (1.59–2.14) but decreased at lower altitudes in French Guiana (0.57–0.78). Pielou's evenness index for species ($J_T$) was relatively constant across altitude classes in Peru (0.79–0.90), whereas it declined at low-altitude sites in French Guiana (0.70-0.76). For mimicry rings, evenness ($J_{MR}$) tended to be slightly higher at low-altitude sites in Peru (0.86–0.87) but showed a marked decrease in one lowland site of French Guiana (0.42).

The NMDS (Non-metric multidimensional scaling) provides a visual representation of taxonomic and mimetic community structure across altitudinal classes. The ordination plots reveal clear separation between low- and high-altitude communities, while mid-altitude communities appear less distinctly clustered (Fig. S5). These patterns are consistent with the differences detected by the PERMANOVA analyses.

The weighted LMMs, which account for the standard deviation of the null model, reveal a significant relationship between the net relatedness index (NRI), which captures the overall structuring pattern across the entire phylogenetic tree, and altitude class (random effects: region variance = 4.66e-10, residual variance = 0.74). Specifically, communities at high altitude exhibited significantly higher NRI values than those at both low and mid-altitudes (high vs. low: estimate = $2.35 \pm 0.80$, $p = 0.05$; high vs. mid: estimate = $2.61 \pm 0.79$, $p = 0.03$). No significant difference was found between low and mid-altitudes (estimate = $0.27 \pm 0.88$, $p = 0.95$) and they showed no significant deviation from randomness (low altitude: $\beta = -0.65 \pm 0.62$, $p = 0.3$; mid-altitude: $\beta = -0.27 \pm 0.87$, $p = 0.8$ –Table S3 & Figure S6). However, for the nearest taxon index (NTI), the effect of altitude class was no significant (random effects: region variance = 1.26e-10, residual variance = 0.39; all $p > 0.2$; see Fig. S6). Hence, high altitude sites with positive NRI and near-zero NTI, are indicative of large-scale phylogenetic clustering with random structure within clades. While mid-altitude sites and low altitude sites, with near 0 NRI and NTI, suggest phylogenetic dispersion close to random.

PERMANOVA showed that altitude was a key driver of phylogenetic β-diversity, explaining 18% of the total variance ($R^2 = 0.18$, $F = 2.30$, $df = 2,6$, $p = 0.03$). Pairwise comparisons revealed significant differences between high and low altitudes ($R^2 = 0.17$, $F = 3.18$, $df = 1,4$, $p = 0.03$), and low and mid-altitudes ($R^2 = 0.15$, $F = 2.42$, $df = 1,4$, $p = 0.04$), whereas differences between high and mid-altitudes were not significant ($p > 0.05$). These results support the existence of phylogenetic differences between low and high altitudes, regardless of region. Moreover, a significant interaction between dissimilarity and region was found (β_nestedness × Region Peru =

–1.65 ± 0.38, $t = –4.34$, $p < 0.001$), indicating that the difference between nestedness and turnover was significantly greater in Peru than in French Guiana. In French Guiana, the contributions of turnover and nestedness to dissimilarity did not differ significantly ($\beta\_nestedness = 0.09 \pm 0.24$, $p = 0.706$), whereas in Peru, nestedness was substantially lower than turnover (difference = –1.56 ± 0.18, $p < 0.001$; Fig. S7).

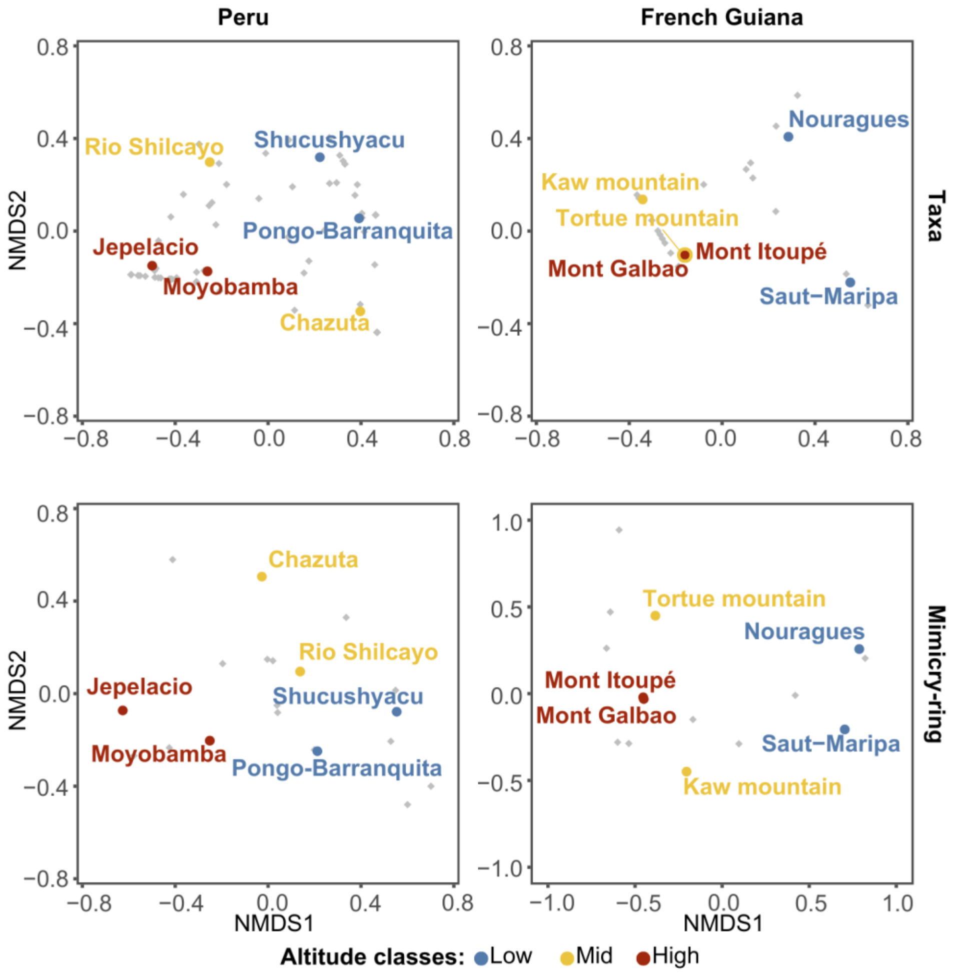


**Figure S5.** Non-metric multidimensional scaling (NMDS) plots based on Bray-Curtis distances for communities of taxa (top) and mimicry rings (bottom). Results are shown for Peru (left) and French Guiana (right), with sites colour-coded by altitude class. Stress values: taxa NMDS – 0 (Peru)*, 4.7e-05 (French Guiana); mimicry NMDS – 0.03 (Peru), 0.03 (French Guiana). *Stress = 0 may reflect low community dissimilarity or limited data structure rather than a perfect ordination. Communities from similar altitude classes cluster more closely than those from different elevations, indicating altitudinal segregation of community composition in both regions.

**Table S3.** Mean NRI and NTI values (± standard deviation) per study site. Positive values indicate phylogenetic clustering, negative values indicate phylogenetic overdispersion. Asterisks (*) indicate that values are significantly different from zero ($p$ < 0.05).

| Region | Sites | NRI | NTI |
|---|---|---|---|
| Peru | Jepelacio (H - 1123m) | 2.3 ± 1.3* | 0.3 ± 1.9 |
| | Moyobamba (H - 955m) | 2.3 ± 1.1* | 0.1 ± 1.6 |
| | Rio shilcayo basin (M - 444m) | -1.5 ± 1.9 | 0.9 ± 3.0 |
| | Chazuta (M - 355m) | -0.05 ±1.7 | -1.3 ± 2.73 |
| | Pongo-Barrenquita road (L - 191m) | -0.5 ± 1.3 | 0.6 ± 2.0 |
| | Shucushyacu (L - 165m) | -0.7 ± 1.2 | 0.6 ± 1.8 |
| French Guiana | Mont Itoupé (H - 826m) | -0.4 ± 1.4 | -1.2 ± 2.7 |
| | Mont Galbao (H - 730m) | 2.6 ± 1.4* | -0.3 ± 3.1 |
| | Tortue Mountain (M - 483m) | -1.9 ± 2.1 | -1.0 ± 4.4 |
| | Kaw Mountain (M - 284m) | -0.6 ± 1.5 | -0.2 ± 5.0 |
| | Nouragues station (L - 54m) | - 1.3 ± 4.6 | 0.2 ± 6.28 |
| | Saut-Maripa (L - 30m) | 0.2 ± 8.1 | 1.3 ± 9.6 |

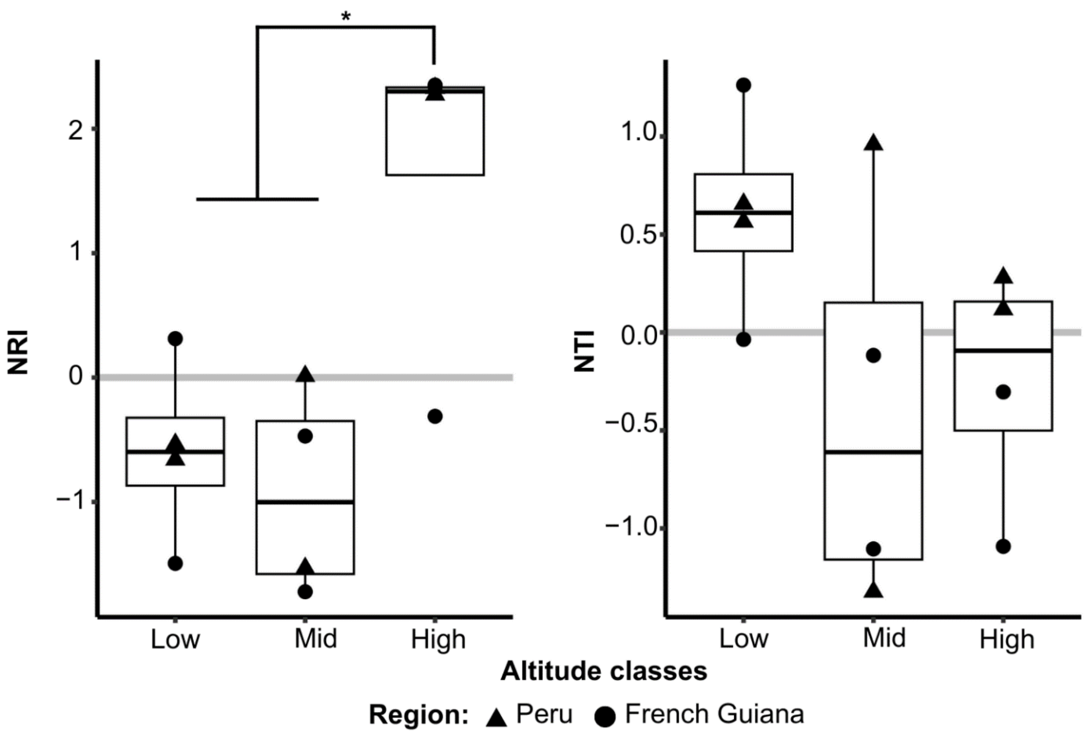


**Figure S6.** Net relatedness index (NRI) and nearest taxon index (NTI) as a function of altitude classes, with shape representing regions. Asterisk (*) indicate significant differences between altitude classes based on weighted linear mixed models ($p < 0.05$). Stronger phylogenetic clustering occurs at higher elevations in both Peru and French Guiana.

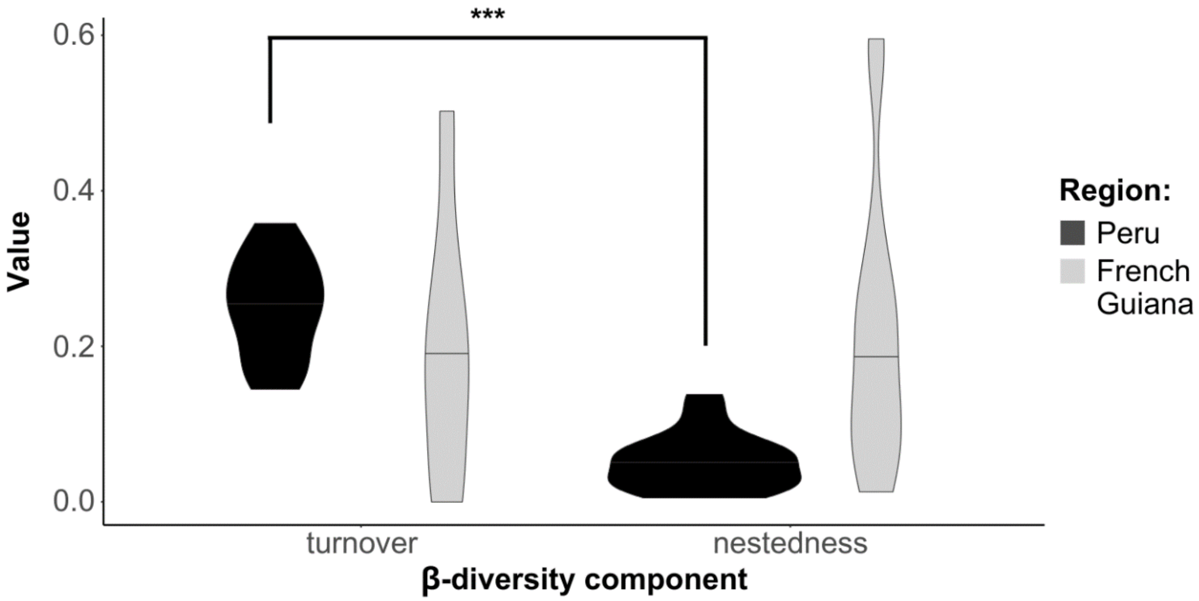


**Figure S7.** Rates of turnover and nestedness between sites for each region, with dark grey representing Peru and light grey representing French Guiana. Asterisks (***) indicate significant differences between β-diversity components based on a generalized linear model ($p < 0.001$). Phylogenetic β-diversity is predominantly driven by turnover rather than nestedness, particularly in Peru where turnover is significantly higher.

*The role of altitude in habitat filtering*

*Climatic variation and elevation: both temperature and precipitation decline with altitude*

GLMs with Gamma distributions were used to assess the relationship between altitude and abiotic factors, specifically temperature and precipitation, for both Peru and French Guiana. The analysis of temperature revealed a significant negative relationship with altitude in both Peru and French Guiana, both for daily temperature during sampling (Peru : $\beta = -0.0002$, $p = 0.03$; French Guiana : $\beta = -0.0001$, $p = 0.007$) and annual mean temperature (Peru : $\beta = -0.0002$, $p < 0.001$; French Guiana : $\beta = -0.0001$, $p < 0.001$), suggesting that higher altitudes are associated with lower temperatures in both regions. The models for precipitation in Peru and French Guiana indicated no significant relationship with altitude for precipitation during sampling (Peru: $\beta = -0.0008$, $p = 0.31$; French Guiana: $\beta = -0.0026$, $p = 0.1$) and a significant negative relationship with altitude for annual precipitation (Peru: $\beta = -0.0005$, $p = 0.025$; French Guiana: $\beta = -0.0003$, $p = 0.025$), indicating that higher altitudes tend to experience lower amounts of precipitation.

*Ithomiini assemblages are not shaped by seasonality but remain structured by altitude*

For the subset of sites in French Guiana sampled in both the rain and dry seasons, altitude class remained a significant factor influencing taxa community composition (df = 2,8, F = 3.53, $R^2 = 0.59$, $p = 0.007$), whereas season (df = 1,8, F = 0.57, $R^2 = 0.05$, $p = 0.80$) was not significant. The interaction between season and altitude class was also non-significant (df = 2,8, F = 0.63, $R^2 = 0.11$, $p = 0.90$) and was not interpreted further due to limited replication. For low-altitude sites, we observed a slight increase in species richness during the rainy season compared to the dry season (Fig. S8). In contrast, species richness remained stable across seasons at mid and high altitudes.

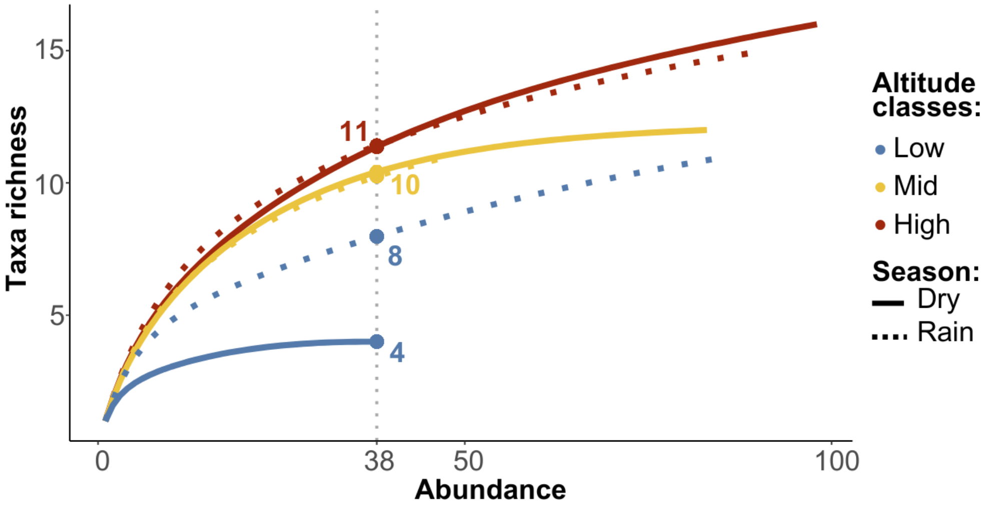


**Figure S8.** Rarefaction curves showing taxa richness in French Guiana as a function of abundance across altitude classes: low altitude (blue), mid-altitude (yellow), and high altitude (red). Solid lines represent the dry season, while dotted lines represent the rain season. The vertical dotted line indicates the minimum sample abundance among sites at which all sites and seasons can be compared. Limited seasonal variation in taxa richness is observed at mid and high elevations, whereas low-altitude sites exhibit higher richness during the rain season.

*Predation pressure across elevation: no altitudinal pattern but a potential driver of diversity and mimicry-based mutualism*

To examine the impact of altitude classes on predation rates, a generalized linear mixed model (GLMM) was used with a binomial distribution, with random effects attributed to different regions. Although the relationship between altitude and predation rate was not statistically significant, a trend toward increased predation at higher elevations was observed (Intercept(low): $\beta = -3.7983 \pm 0.4$, $z = -8.6$, $p < 2e\text{-}16$; mid: $\beta = 0.27 \pm 0.3$, $z = 0.9$, $p = 0.4$; high: $\beta = 0.3 \pm 0.3$, $z = 1$, $p = 0.3$), particularly in the Peruvian sites (Fig. S9). However, a significant difference was observed between the two regions, with this factor explaining 30% of the model's variance; indeed, predation rates were highest for Peru, regardless of altitude (Variance = 0.28, SD = 0.53).

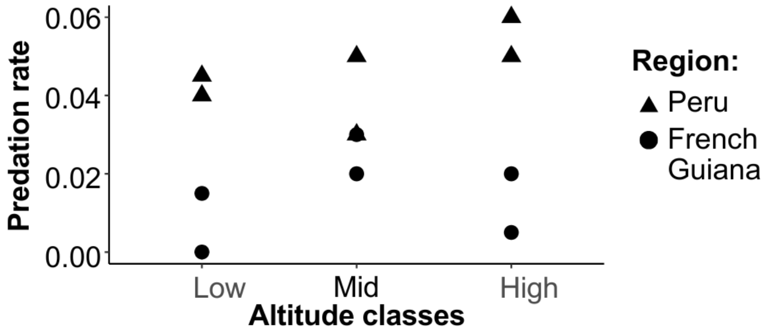


**Figure S9.** Predation rates by altitude class, with triangles indicating sites in Peru (N=6 sites) and circles indicating sites in French Guiana (N=6 sites). Predation pressure does not vary systematically along the altitudinal gradient in either region.

GLMMs (with a quasi-Poisson distribution) were used to explore the impact of predation rates on species and mimicry ring richness, while accounting for regions and revealed a significant positive effect of predation rate on species richness ($\beta = 20.68 \pm 5.24$, $t = 3.95$, $p = 0.003$; Fig. 2a), indicating that higher predation rates are correlated with increased biodiversity within the studied localities. Similarly, a positive correlation was found between predation rate and mimicry-ring richness ($\beta = 14.96 \pm 3.45$, $t = 4.33$, $p = 0.002$; Fig. 2b). Additionally, a positive trend was found between predation rate and the average number of taxa per mimicry ring ($\beta = 6.16 \pm 2.90$, $t = 2.13$, $p = 0.062$) and a significant positive correlation between the median number of taxa per mimicry-rings and predation rate ($\beta = 18.13 \pm 3.83$, $t = 4.72$, $p = 0.001$; Fig. 2c). The random effect of region remained negligible for all tests (SD comprised between 7.49e-07 and 1.93e-06), indicating similar patterns in Peru and French Guiana.

*Diversity begets diversity: host plant richness increases with elevation and seems to promote Ithomiini diversity*

The performance of the ensemble species distribution models (SSDMs) for Ithomiini butterflies and their host plants (Solanaceae and Gesneriaceae) was assessed using a range of algorithms (Table S4). Evaluation metrics included the area under the curve (AUC), omission rate, sensitivity, specificity, proportion of correct classifications, Cohen's Kappa, and calibration score.

**Table S4.** Evaluation of species distribution models (SSDMs). Evaluation using different modelling algorithms: artificial neural networks (ANN), classification tree analysis (CTA), generalized additive models (GAM), generalized boosted regression models (GBM), generalized linear models (GLM), multivariate adaptive regression splines (MARS), maximum entropy (MXT), random forest (RF), and support vector machines (SVM).

| Group | Model | Threshold | AUC | Omission rate | Sensitivity | Specificity | Prop correct | Kappa | Calibration | Kept model |
|---|---|---|---|---|---|---|---|---|---|---|
| Ithomiini | ANN | 0.72 | 0.75 | 0.27 | 0.86 | 0.61 | 0.73 | 0.46 | 0.82 | 1.33 |
| | CTA | 0.94 | 0.83 | 0.17 | 0.83 | 0.82 | 0.83 | 0.65 | 0.45 | 8.19 |
| | GAM | 0.09 | 0.90 | 0.16 | 0.84 | 0.84 | 0.84 | 0.29 | 0.85 | 9.38 |
| | GBM | 0.04 | 0.90 | 0.17 | 0.83 | 0.83 | 0.83 | 0.66 | -0.50 | 9.19 |
| | GLM | 0.10 | 0.87 | 0.19 | 0.81 | 0.81 | 0.81 | 0.26 | 0.83 | 8.48 |
| | MARS | 0.05 | 0.87 | 0.17 | 0.79 | 0.84 | 0.83 | 0.27 | 0.84 | 8.76 |
| | MXT | 0.17 | 0.90 | 0.50 | 0.89 | 0.50 | 0.50 | 0.01 | 0.53 | 9.2 |
| | RF | 0.53 | 0.92 | 0.15 | 0.85 | 0.85 | 0.85 | 0.69 | 0.53 | 9.52 |
| | SVM | 0.47 | 0.90 | 0.17 | 0.83 | 0.82 | 0.83 | 0.65 | 0.44 | 9.33 |
| Host plants | ANN | 0.74 | 0.78 | 0.27 | 0.83 | 0.64 | 0.73 | 0.47 | 0.58 | 1.22 |
| | CTA | 0.92 | 0.81 | 0.20 | 0.82 | 0.78 | 0.80 | 0.60 | 0.39 | 6.74 |
| | GAM | 0.08 | 0.85 | 0.22 | 0.78 | 0.78 | 0.78 | 0.20 | 0.84 | 9.21 |
| | GBM | 0.13 | 0.86 | 0.21 | 0.79 | 0.78 | 0.79 | 0.57 | -0.48 | 8.70 |
| | GLM | 0.08 | 0.83 | 0.25 | 0.76 | 0.75 | 0.75 | 0.17 | 0.85 | 8.80 |
| | MARS | 0.05 | 0.82 | 0.22 | 0.74 | 0.79 | 0.78 | 0.20 | 0.79 | 7.51 |
| | MXT | 0.36 | 0.85 | 0.50 | 0.84 | 0.50 | 0.50 | 0.01 | 0.47 | 9 |
| | RF | 0.55 | 0.89 | 0.18 | 0.82 | 0.82 | 0.82 | 0.63 | 0.52 | 9.14 |
| | SVM | 0.54 | 0.86 | 0.21 | 0.79 | 0.79 | 0.79 | 0.58 | 0.41 | 8.49 |

407

Furthermore, the GAM analysis revealed a strong and significant relationship between host plant species richness and Ithomiini species richness across altitudinal classes (low : 0-250m, mid : 250-500m, high : 500-1200m; see Fig. 7). The model explained 87.6% of the deviance, with an adjusted $R^2$ value of 0.87, indicating a highly robust model fit. The F-statistic was significant (edf = 10.63, Ref.df = 11, F = 61053, $p < 0.001$), indicating a positive correlation between host plant richness and Ithomiini richness.

Although the assumptions of multivariate normality and homogeneity of covariance matrices were not met (Box's M test: $\chi^2 = 328.74$, df = 6, $p < 0.001$), a linear discriminant analysis (LDA) was conducted as an exploratory approach to assess whether Ithomiini taxa richness and host plant species richness differed between altitude classes. The projection revealed a clear separation, particularly between low- and high-altitude sites (Fig. S10). The first linear discriminant (LD1), which explained 98.3% of the variance, was primarily associated with Ithomiini richness (coefficient = 0.85), while the second axis (LD2) contributed minimally (1.7%). The overall classification accuracy was 75.8%, with most misclassifications occurring between adjacent altitude classes (low vs. mid, mid vs. high).

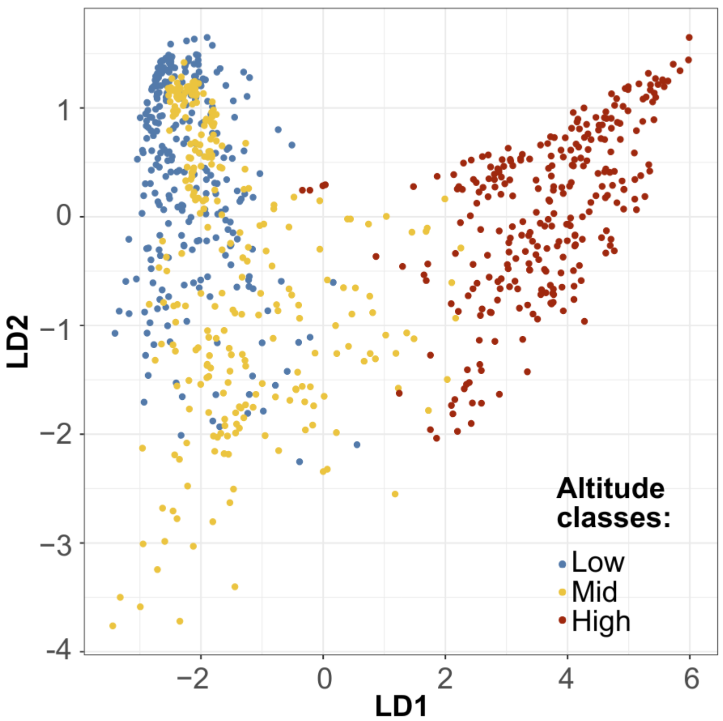


**Figure S10.** Linear discriminant analysis (LDA) showing the separation of altitude classes based on Ithomiini taxa richness and host plant species richness in French Guiana. The underlying data consist of 343,957 observations modelled using the SSDM package; for visualization purposes, a subset of 277 randomly sampled points per altitude class was used. Altitude categories are represented by colour. Combined variation in butterfly and host plant richness discriminates altitude classes, with high-altitude communities clearly separated from low- and mid-elevation assemblages.

*Ecological factors shape Ithomiini assemblages across regions with differing evolutionary time*

To assess the influence of ecological factors on variation in community assemblages, we performed a multivariate distance matrix regression (MDMR) using a set of environmental and biotic predictors. The predictor set included annual mean temperature, annual precipitation, mean forest cover, and predation rate, selected after evaluating multicollinearity using variance inflation factor (VIF) analysis. Mixed MDMR analysis confirmed a significant effect of ecological variables on Ithomiini taxa composition (Statistic = 8.17, DF = 4, $p = 0.006$), independently of regional effects. Among the tested predictors, predation rate showed a strong and significant contribution to community composition (Statistic = 3.63, $p = 0.005$), suggesting that predator-mediated selection plays a key role in shaping assemblages. In contrast, annual mean temperature showed only a marginal trend (Statistic = 2.18, $p = 0.051$), and annual precipitation and forest cover had no detectable effect on community dissimilarity (Precipitation: Statistic = 0.95, $p = 0.40$; Forest cover: Statistic = 1.49, $p = 0.20$).
